# BayesTME: A unified statistical framework for spatial transcriptomics

**DOI:** 10.1101/2022.07.08.499377

**Authors:** Haoran Zhang, Miranda V. Hunter, Jacqueline Chou, Jeffrey F. Quinn, Mingyuan Zhou, Richard White, Wesley Tansey

## Abstract

Spatial variation in cellular phenotypes underlies heterogeneity in immune recognition and response to therapy in cancer and many other diseases. Spatial transcriptomics (ST) holds the potential to quantify such variation, but existing analysis methods address only a small part of the analysis challenge, such as spot deconvolution or spatial differential expression. We present BayesTME, an end-to-end Bayesian method for analyzing spatial transcriptomics data. BayesTME unifies several previously distinct analysis goals under a single, holistic generative model. This unified approach enables BayesTME to (i) be entirely reference-free without any need for paired scRNA-seq, (ii) outperform a large suite of methods in quantitative benchmarks, and (iii) uncover a new type of ST signal: spatial differential expression within individual cell types. To achieve the latter, BayesTME models each phenotype as spatially adaptive and discovers statistically significant spatial patterns amongst coordinated subsets of genes within phenotypes, which we term spatial transcriptional programs. On human and zebrafish melanoma tissues, BayesTME identifies spatial transcriptional programs that capture fundamental biological phenomena like bilateral symmetry, differential expression between interior and surface tumor cells, and tumor-associated fibroblast and macrophage reprogramming. Our results demonstrate BayesTME’s power in unlocking a new level of insight from spatial transcriptomics data and fostering a deeper understanding of the spatial architecture of the tumor microenvironment. BayesTME is open source and publicly available (https://github.com/tansey-lab/bayestme).

## Main

The tumor microenvironment (TME) is composed of a heterogeneous mixture of cell phenotypes, subtypes, and spatial structures. The composition of the TME impacts disease progression and therapeutic response. For instance, the composition of immune cells in the tumor microenvironment is a determinant of response to immunotherapy (IO)^1^. More recent work suggests that it is not cellular composition but rather the *spatial organization* of the microenvironment that determines IO response^2,3,4,5^. Spatially-unaware approaches, such as single-cell RNA and DNA sequencing (scRNA-seq and scDNA-seq), are able to capture the presence and abundance of different cell types and phenotypes (hereon referred to as simply types)^6^, but are unable to characterize their spatial organization. Spatial measurements and spatial modeling of the tumor microenvironment *in situ* present an opportunity to fully uncover and understand the role that spatial structure plays in determining disease progression and therapeutic response.

Spatial transcriptomics (ST) technologies, such as Visium^7^, HDST^8^, and Slide-seq^9^ enable biologists to measure spatially-resolved gene expression levels at thousands of spots in an individual tissue. Each tissue is divided into a grid or lattice of spots, with each spot in the grid typically 50–100*µm* wide, typically covering 10–60 cells. The tissue is permeabilized to release mRNAs to capturing probes with spot-specific barcodes. Bulk RNA-seq is then run on the captured mRNAs tagged with spatial barcodes. The result is a high-dimensional, spatially-localized gene expression count vector for each spot, representing an aggregate measurement of the gene expression of the cells in the spot.

Modeling spot-wise aggregate measurements is challenging as it requires disentangling at least four sources of spatial variation present in the raw signal. First, technical error, also known as spot bleeding, causes mRNAs to bleed to remote spots and contaminates the raw spatial signal. Second, variation in cell counts changes the absolute number of unique molecular identifiers (UMIs) per spot. Since UMI counts scale with the number of cells in each spot, conventional pre-processing methods like log-normalization break this linear relationship. Third, differences in the cell type proportions in each spot conflate signal strength with cell type prevalence. This complicates analysis as it necessitates performing a difficult deconvolution of each spot into its constituent cell type composition. These three sources of variation obscure the fourth, namely the spatial variation in gene expression within each cell type in response to the microenvironment. Teasing out these different sources of spatial variation in ST data is necessary to obtain a full understanding of the spatial architecture of the tumor microenvironment.

Several methods have been developed that specialize in a subset of these four sources of spatial variation. SpotClean^10^ corrects spot bleeding by fitting an isotropic Gaussian model to raw UMI counts in order to map them back to their most likely original location. Spatial clustering methods^11,12,13^ fuse spots together to effectively capture regions of constant cell type proportion with varying cell counts. Spot deconvolution methods^14,15,16^ separate the aggregate signals into independent component signals with each attributable to a different cell type. Spatial differential expression methods^17,18^ assess the aggregate spot signal to detect regions where individual genes or gene sets follow a spatial pattern. While each of these methods has moved the field of ST analysis forward, they each have shortcomings such as making incorrect parametric assumptions, requiring perfect reference scRNA-seq data, or only capturing aggregate signals rather than phenotype-specific ones.

Notably, existing methods assume cells of a given type have a static distribution of gene expression. This assumption is at conflict with the biological knowledge that cells change their behavior in response to their local microenvironment under mechanisms including proliferation, invasion, and drug resistance^19^. The microenvironment regulates cell behavior and therefore alters gene expression profiles of specific cell phenotypes^20^. These microenvironmental influences are particularly relevant in disease contexts. As an example, the microenvironment affects each phase of cancer progression and invasion-metastasis cascade^21^. Chronic inflammation is able to induce tumor initiation, malignant conversion, and invasion^22^. Recent research also shows cancer cells in the interior of a tumor behave differently than cancer cells at the interface with healthy cells^23^. Existing methods are unable to accurately capture spatial expression variation within cell types and thus modeling ST data to understand the spatial structure of transcriptomic diversity in each cell type remains an important open problem.

In this paper, we present BayesTME, a holistic Bayesian approach to end-to-end modeling of ST data that goes beyond existing techniques and captures spatial differential expression within cell types. BayesTME uses a single generative model to capture the multiscale and multifaceted spatial signals in ST data. At the highest level, BayesTME models the global pattern of spatial technical error present in raw ST data. As we demonstrate, ST data contain technical error that is anisotropic, with UMIs bleeding toward a specific direction in each sample. At the intermediate level, BayesTME places spatial fusion priors between spots, adaptively fusing tissue regions together to reveal cellular community structure. This also enables BayesTME to pool statistical strength across spots, enabling it to perform spot deconvolution without single-cell RNA-seq reference. Graph smoothing priors are simultaneously used to capture the spatial heterogeneity of within-phenotype gene expression. These priors enable BayesTME to discover spatial transcriptional programs (STPs), coordinated spatial gene expression patterns among groups of genes within a phenotype. Through an efficient empirical Bayes inference procedure, BayesTME infers all of the latent variables in the generative model with full quantification of uncertainty. Thus, BayesTME provides statistical control of the false discovery rate for marker genes, cell counts, expression profiles, and spatial transcriptional programs. Figure 1 provides an overview of the BayesTME computational workflow (top) and outputs (bottom).

**Figure 1:**
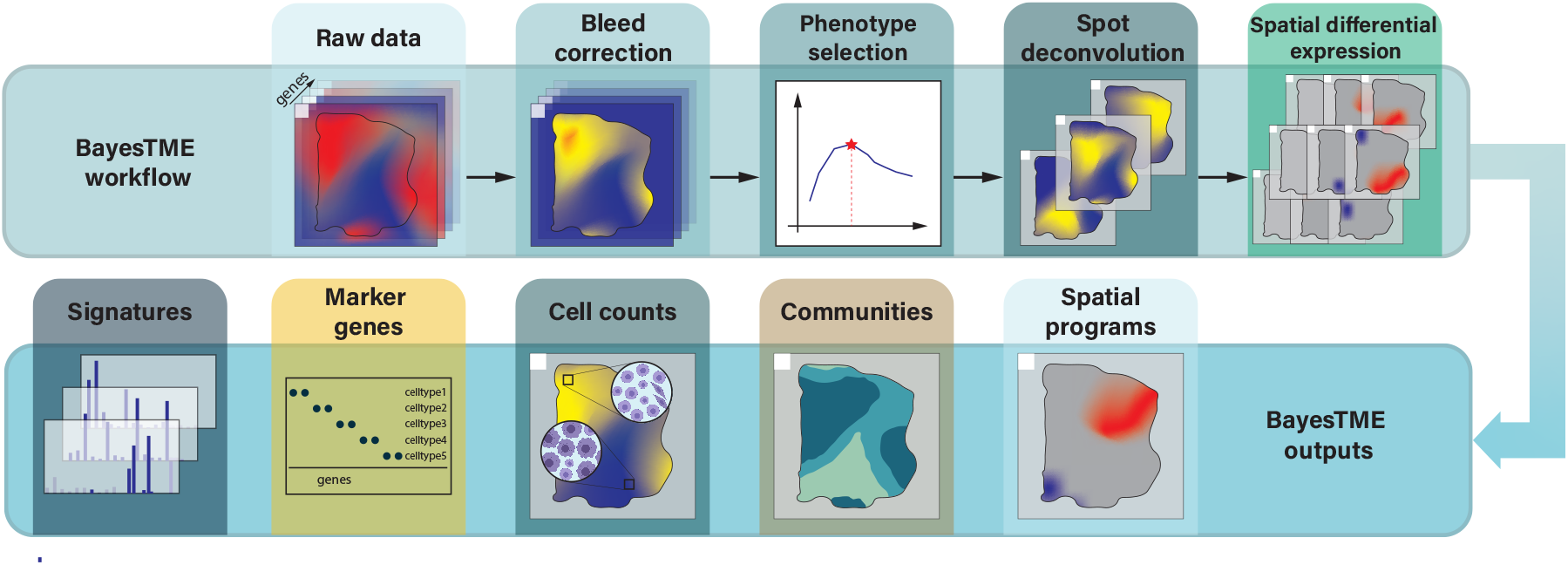
The BayesTME computational workflow and outputs. **Top:** BayesTME first corrects technical errors (spot bleeding) in the raw ST data by probabilistically mapping reads to their most likely original location in the tissue. An unbiased spatial cross-validation routine is then run to select the optimal number of distinct cell phenotypes. The cell phenotype count is then fixed and a reference-free spot deconvolution is run to simultaneously recover the cell phenotypes and their counts at each spot. Finally, the deconvolution model is augmented with a spatially-adaptive phenotype model to infer phenotype-specific spatial variations. **Bottom:** The final output of the complete BayesTME pipeline is the inferred cell phenotype expression signatures, the top marker genes that maximally distinguish phenotypes, the posterior distribution over the discrete cell counts of each type in each spot, the segmented tissue partitioned into cellular communities, and the spatial transcriptional programs discovered for each phenotype.

BayesTME outperforms existing methods on benchmarks for bleed correction, cell type identification, spot deconvolution, cellular community segmentation, and within-phenotype spatial gene expression. We demonstrate that existing methods based on aggregate spatial differentiation are unable to detect within-phenotype variation due to spatial variation in cell type proportions. In contrast, BayesTME identifies spatial transcriptional programs with high power while maintaining tight control over the false discovery rate on the reported spatially-varying genes in each cell type. On real tissues from human melanoma and a zebrafish melanoma model, BayesTME identifies spatial programs that capture core biological concepts like bilateral symmetry and differential expression between the surface and interior tumor cells. BayesTME is open source^1^, does not require reference scRNA-seq, and all hyperparameters are auto-tuned without the need for any manual user input.

## Results

### A holistic generative model for spatial transcriptomics

BayesTME models spatial variation at multiple scales in ST data using a single hierarchical probabilistic model. At the top-level, spot bleeding is modeled via a semi-parametric spatial contamination function. This bleeding model allows for any arbitrary spot bleeding process to be modeled, under the constraint that UMIs are less likely to bleed to spots that are farther away. By leveraging the non-tissue regions as negative controls (i.e. spots where the UMI count should be zero), BayesTME learns this function and then inverts it to estimate the true UMI counts for each in-tissue spot.

At the spot level, BayesTME models true UMI counts in each spot using a carefully specified negative binomial distribution. The spot convolution effects due to cell aggregation in each spot are captured in the rate parameter. This ensures that a linear increase in the number of a particular cell type yields a linear increase in the UMIs from that cell type. The success probability parameter in the negative binomial likelihood is used to capture spatial variation *within each cell type*. These latter spatial parameters allow cell types to up- or down-regulate genes in each spot, enabling BayesTME to capture dynamic phenotypic behavior at spatially localized regions in the TME. This careful separation enables BayesTME to capture within phenotype spatial variation of gene expression, a more nuanced signal than currently recoverable by existing methods. Further, the uncertainty quantification provided by posterior inference enables BayesTME to detect significantly varying genes in each cell type with control of the false discovery rate.

Hierarchical priors in BayesTME encode heavy-tailed Bayesian variants^24,25^ of the graph-fused group lasso prior^26^ and the graph trend filtering prior^27^. The fused lasso prior enforces that the prior probability distribution over cell types follows a piecewise constant spatial function, encoding the biological knowledge that groups of cell phenotypes form spatially contiguous communities. The graph trend filtering prior allows gene expression to vary within cell types, encoding the biological knowledge that cells execute gene sets in a coordinated fashion, known as transcriptional programs. Spatial transcriptional programs extend this concept by identifying and quantifying the activation level of different programs in space. Identification of the BayesTME parameter values is achieved through a novel empirical Bayes inference algorithm that enables Bayesian quantification of uncertainty over each parameter of interest in the decontaminated data. See the Methods for the detailed hierarchical specification of the generative model and for details on parameter estimation.

### BayesTME accurately corrects previously-unreported directional spot bleeding in ST data

Plots of raw UMI counts in real ST data (Figure 2a-c) show the UMI signal bleeds to background spots with a gradient of intensity. These plots also suggest, unlike the Gaussian assumption in previous preprocessing methods^10^, or the uniform background noise model in other models^16^, bleeding error varies in magnitude in different directions. Such phenomena may be the result of cell-free DNA from dead cells, mRNA binding capacity limitation of spatial barcodes, or technical artifacts of tissue permeabilization.

**Figure 2:**
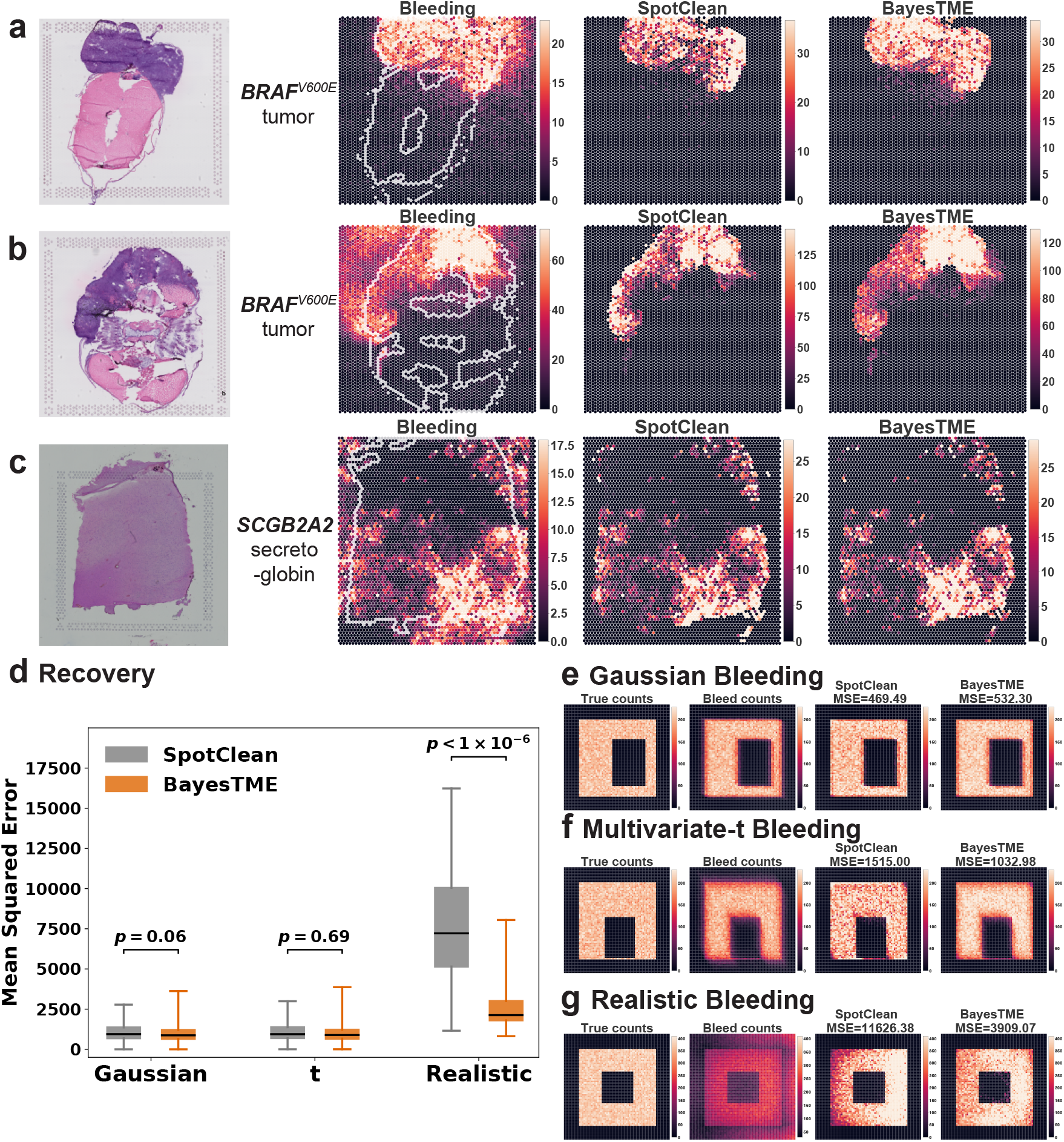
BayesTME recovers UMI reads from bleeding contamination and preserves the spatial pattern of interest. (**a-c**.) Bleed correction of selected marker genes in (**a-b**.) two zebrafish melanoma model samples and (**c**.) a human dorsolateral prefrontal cortex sample^28^, with comparison to SpotClean. Bleeding patterns consistently show directional, anisotropic skew towards one corner. SpotClean UMI corrections are therefore expected to be biased towards the tissue boundary whereas BayesTME is more diffuse and better recapitulates the true signal. (**d**.) BayesTME performs similarly to SpotClean when the bleeding pattern is isotropic and not skewed (e.g. Gaussian or Student’s t); BayesTME substantially outperforms SpotClean when bleeding skews UMIs toward one direction as observed in real tissues. (**e-g**.) Examples of simulated bleeding patterns showing how BayesTME is able to learn and correct for the direction of the bleeding pattern.

BayesTME corrects bleeding while preserving the true signal. To do this, BayesTME learns a semi-parametric anisotropic bleeding model to correct directional ST bleed and map UMIs to their most likely origin in the tissue. The BayesTME correction only assumes that UMI bleeding decays monotonically as a function of distance. Non-tissue regions are leveraged by BayesTME as a form of negative control, enabling the method to identify the underlying spatial error function from the data via a maximum likelihood estimation procedure.

To evaluate the performance of the BayesTME bleed correction, we constructed synthetic datasets simulating three different bleeding mechanisms: Gaussian, heavy-tailed multivariate-t, and realistic (anisotropic) direction-biased bleeding (Figure 2e-g). The last simulation was constructed to resemble real ST data, with bias towards a specific corner of the slide. We compared BayesTME with SpotClean^10^ (Figure 2d), an existing ST error correction technique that assumes Gaussian technical error. While both methods perform comparably in Gaussian (*µ*_*MSE,SpotClean*_ = 1170.08, *µ*_*MSE,BayesTME*_ = 1263.66, *p*-value = 0.06) and multivariate-t (*µ*_*MSE,SpotClean*_ = 1210.06, *µ*_*MSE,BayesTME*_ = 1305.31, *p*-value = 0.69) bleeding scenarios, BayesTME significantly outperformed SpotClean in the realistic bleeding scenario (*µ*_*MSE,SpotClean*_ = 10437.48, *µ*_*MSE,BayesTME*_ = 3048.92, *p*-value = 1.87 × 10^−301^).

We found that cell typing and deconvolution were robust to this spatial error. However, bleed correction was critical to preventing genes from falsely registering as spatially varying in real ST data. These results suggest that ST experimental workflows should take care to allow ample non-tissue space in each direction of the slide. If the tissue section exceeds the fiducial markers substantially in a given direction, the technical error function will be statistically unidentifiable. In such cases, it will be impossible to distinguish technical error from true spatial variation, potentially leading to false conclusions when assessing spatially-varying gene expression within-phenotypes.

### BayesTME outperforms a suite of existing methods for phenotype inference, spot deconvolution, and tissue segmentation

We benchmarked BayesTME against other methods: BayesSpace^11^, cell2location^16^, DestVI^15^, CARD^29^, RCTD^30^, STdeconvolve^14^, stLearn^12^, and Giotto^13^ on simulated data based on real single-cell RNA sequencing (scRNA) data. We randomly sampled *K*\* cell types from a previously-clustered scRNA dataset^16^; we conducted experiments for *K*\* from 3 to 8. For a given *K*\*, we constructed spatial layouts consisting of 25 cellular communities, defined as spatially-contiguous regions of homogeneous mixtures of cell types. We randomly generated the total cell number for each spot with cellular-community-specific priors. After dividing the total cell number into *K*\* cell types, we randomly sampled cells from the scRNA data of the selected cell types and mapped them on top of the spot pattern from a human melanoma tissue sample^31^; see the Supplement for details. We compared the performance of BayesTME to the above existing methods on selecting the correct number of cell phenotypes, deconvolving spots, segmenting tissues into spatial communities, and detecting groups of spatially-varying genes within phenotypes. As we demonstrate, BayesTME outperformed existing methods across all benchmark tasks.

### BayesTME accurately identifies the correct number of cell phenotypes and each phenotype expression signature

A core modeling task in ST analysis is deconvolution of the spots into their constituent cell phenotype proportions. Most existing methods require a scRNA-seq reference for deconvolution and cell type mapping. As has been noted^16^, these methods may be brittle when a cell type is missing from the reference. This vulnerability is particularly problematic in cancer where many subclones may exist and non-overlapping sets of subclones occur between different tissue samples. BayesTME learns the cell phenotypes–both the number of types and their signatures–directly from ST data without the need for scRNA-seq. Thus, BayesTME is robust to the natural spatial heterogeneity of phenotypes in cancer and other disease tissues. To evaluate the robustness and performance of BayesTME, we compared it to both an existing reference-free method and to existing reference-based methods with different degrees of scRNA missingness. To focus purely on the deconvolution and reference-free capabilities of BayesTME, our simulations did not apply any spot bleeding.

There are two tunable hyperparameters in BayesTME: *K*, the number of cell types, and *λ*, the global degree of smoothness. BayesTME uses a spatial cross-validation approach to automatically select both variables without the need for user input. The cross-validation procedure creates *m* non-overlapping folds each with *κ*% of spots held out; we set *m* = 5 and *κ* = 5%. For each fold, BayesTME enumerates *K* = 2, …, *K*_max_ and *λ* = 10^1^, …, 10^6^; in all of our experiments we set *K*_max_ = 15. For each (*K, λ*), we fit BayesTME on the in-sample data. Graph smoothing priors enable BayesTME to fill-in missing spots during cross-validation. BayesTME uses these imputed posterior estimates to evaluate the likelihood on the held out data. BayesTME integrates out *λ* in order to select *K* then chooses the *λ* value closest to the mean held out likelihood for the chosen *K*; see the Methods for details.

We first evaluated how well the BayesTME recovers the true gene expression profiles of each cell type in each of our *K*\* (true number of cell types) settings. We compared the BayesTME result to STdeconvolve, a reference-free alternative method based on latent Dirichlet allocation^32^ that provides three different approaches to estimating the number of cell types; we picked the closest estimation out of the three candidates that STdeconvolve provided. Reference-based methods assume access to ground truth cell type information from scRNA annotation, making them unavailable for comparison. In each simulation, BayesTME achieved a higher correlation with the true gene expression levels as measured by *r*^2^ (Figure 3a). Further, STdeconvolve over- or underestimated the true number of cell types whereas BayesTME selected the correct number of cell types in each setting (Figure 3c, left).

**Figure 3:**
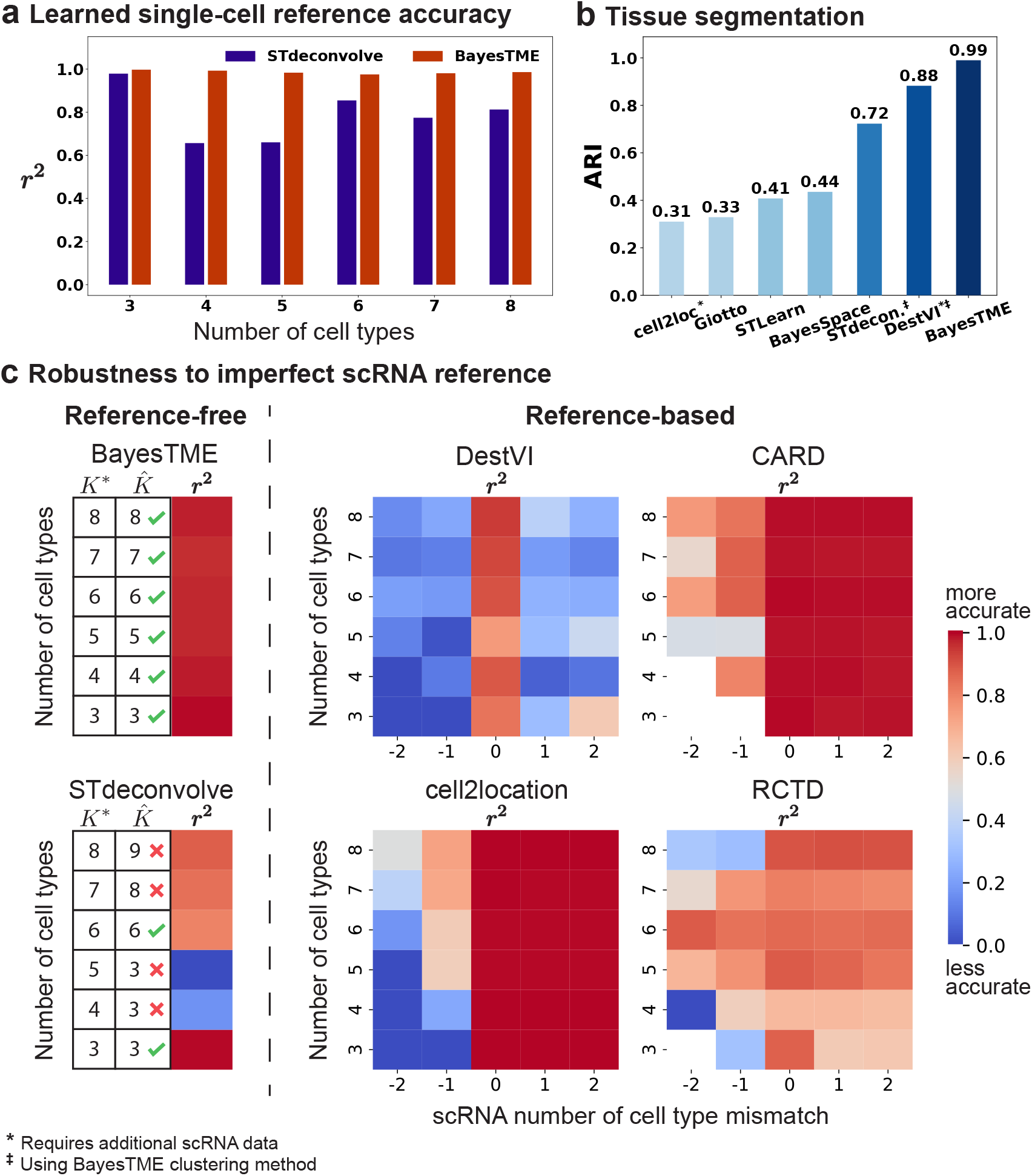
BayesTME performs outperforms existing methods in semi-synthetic benchmarks. (**a**.) BayesTME outperforms the reference-free method STdeconvolve in expression profile inference for each cell type, measured by the coefficient of determination (*r*^2^), for semi-synthetic data with ground truth number of cell types *K*^*∗*^ = 3, 4, 5, 6, 7, 8 (**b**.) BayesTME outperforms all other methods when segmenting the tissue into cellular communities, measured by adjusted Rand index (ARI). (**c**.) BayesTME outperforms existing methods in robustness benchmarks. Reference-based methods are vulnerable to imperfect scRNA reference as demonstrated by the decline in spot deconvolution accuracy; x-axis: reference contains a subset (< 0), exact match (= 0), or superset (> 0) of the true reference. The existing reference-free method is not reliable in picking the correct number of cell types. BayesTME simultaneously detects the optimal number of cell types from the data and accurately deconvolves the spots.

We next evaluated the robustness of reference-based methods DestVI, CARD, cell2location, and RCTD to reference mismatch. We found that while all methods performed well when the reference was perfectly matched, reference mismatch was problematic for all four reference-based methods (Figure 3c, right). Specifically, DestVI and RCTD were sensitive to the reference being a superset of the true number of cell types (x-axis values 1 and 2) and all four were sensitive to missing cell types (x-axis values -1 and -2). By not relying on any reference scRNA-seq, BayesTME retained high accuracy across all simulations (Figure 3c, left).

Finally, we evaluated the ability of different methods to segment the tissue into spatial regions representing cellular communities. In community detection benchmarks, BayesTME (adjusted Rand index^33^, *ARI* = 0.99) surpassed all other currently available alternatives (Figure 3 b), including both spatial clustering (BayesSpace, STLearn, Giotto) and spot deconvolution (cell2location, DestVI, STdeconvolve) methods. For cell2location (*ARI* = 0.31) we used its built-in Leiden clustering; when inserting the BayesTME spatial clustering, cell2location improved to *ARI* = 0.93, suggesting the BayesTME clustering provides an independent benefit even for accurate deconvolution methods.

### BayesTME identifies within-phenotype spatial transcriptional programs with tight control of the false discovery rate

In addition to bleed correction, deconvolution, and cell typing, BayesTME detects gene expression levels of each phenotype that vary in space. To do this, the generative model for BayesTME uses a negative binomial likelihood where spatially-invariant expression levels parameterize the rate and spatially-dependent expression levels parameterize the success rate. Hierarchical spatial shrinkage and clustering priors on the success rate parameters enable BayesTME to discover genes within each phenotype that spatially vary in coordination with other genes. We call these gene sets and spatial patterns *spatial transcriptional programs* (STPs). The STP construction in BayesTME is flexible: it allows for genes to be negatively spatially correlated within the same program, makes no assumption on the shape or pattern of spatial variation, and adaptively discovers how many genes are in each program. After inference, we use the posterior uncertainty to select STPs with control of the Bayesian false discovery rate (see Methods); we set the FDR target to 5% by default.

To benchmark BayesTME, we constructed a simulation dataset with spatial transcriptional programs by randomly sampling cells from the scRNA data following the same fashion as in the previous experiments. We used the spatial layout from a zebrafish melanoma sample as it is a large tissue containing more than 2000 spots, enabling a rich set of spatial patterns to be imprinted. We chose *K*\* = 3 cell types and designed 2 spatial programs for each cell type, where 10 genes were randomly sampled and assigned to each of the STPs (Figure 4c). After selecting these 60 spatial genes, we reordered their sampled reads by the spot intensity of their respective spatial programs to simulate the spatial differentiation while preserving the mean expression. Thus, while the gene expression patterns are spatially informative in these simulations, clustering by scRNA-seq analysis would remain unchanged.

**Figure 4:**
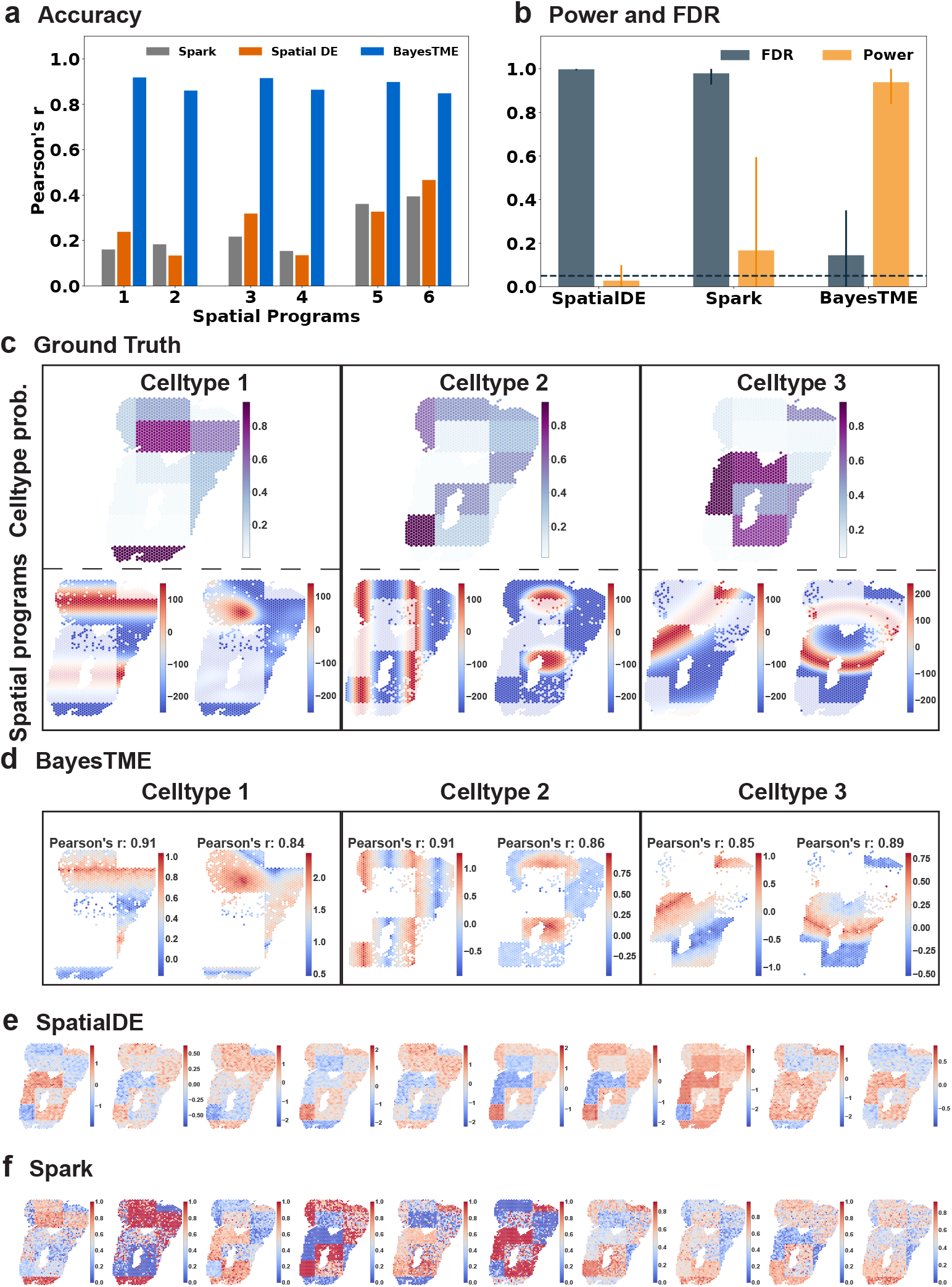
BayesTME discovers spatial transcriptional programs with high power and tight control of the false discovery rate. (**a**.) Accuracy of the closest spatial pattern discovered by each method to the ground truth. (**b**.) True positive rate (orange) and false discovery rate (gray) for each method when predicting which genes belong to each spatially varying pattern; the dashed line is the target (5%) false discovery rate. (**c**.) Ground truth spatial patterns used in the benchmark simulations; top: cell type proportion probabilities; bottom: spatial pattern followed by the genes in each spatial program. (**d**.) Spatial programs found by BayesTME at the 5% FDR level. (**e-f**.) Spatial patterns found by other methods; both SpatialDE and Spark are unable to disentangle phenotype proportions from spatial gene expression within phenotypes.

We benchmarked BayesTME against spatial differential expression methods^18,17^ that enable control of the false discovery rate. BayesTME identified all 6 spatial transcriptional programs with on average 0.88 Pearson’s *r* correlation to the ground truth (Figure 4a,c). In contrast, we found SpatialDE and Spark could only detect phenotype proportion patterns instead of meaningful within-phenotype variation in spatial gene expression (Figure 4d-e). We also evaluated the DestVI spatial expression detection mechanism and found the results to be uncorrelated with the ground truth (see Supplement for details). Quantitatively, BayesTME achieved an average false discovery proportion of 14% where the 95% confidence interval covers the 5% target FDR, and TPR of 94% for selecting spatially varying genes (Figure 4b).

### BayesTME discovers novel spatial programs of immune infiltration and response in human melanoma

We applied BayesTME to a published human melanoma dataset^31^ generated using first generation ST technology, with a spot diameter of 100 *µm* and center-to-center distance between spots of 200 *µm*^34^. *The* selected sample contained visible tumor, stromal, and lymphoid tissues as annotated by a pathologist based on H&E staining (Figure 5a). Despite the relatively low resolution of the data, the cell types identified by BayesTME successfully recapitulated the histology of the tissue (Figure 5b).

**Figure 5:**
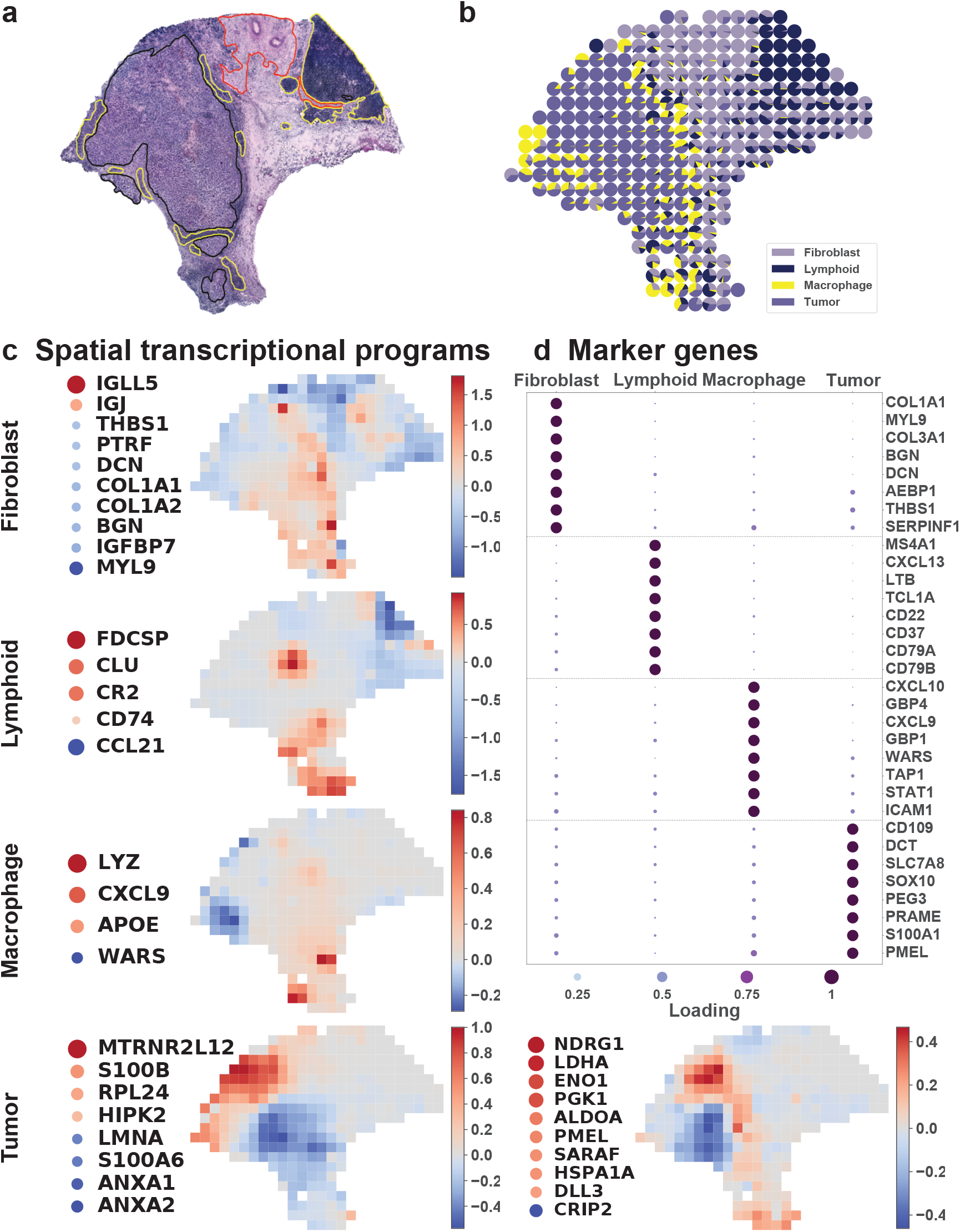
BayesTME discovers novel spatial programs of immune-tumor interaction in human melanoma. (**a**.) Pathologist-annotated H&E slide; yellow: immune cells, red: stroma, black: tumor. (**b**.) BayesTME recovers 4 cell types which map closely to the pathologist annotations. (**c**.) BayesTME recovers 5 spatial programs representing fibroblasts, immune cells, and two programs covering tumor subtypes related to transcription (left) and stress responses (right). (**d**.) Top marker genes selected by BayesTME to describe each cell type.

Five spatial transcriptional programs were identified by BayesTME (Figure 5c). Two programs were tumor-specific, and displayed somewhat distinct expression patterns, suggesting a spatially-segregated pattern of tumor heterogeneity (Figure 5c). As expected, melanoma marker genes such as *PMEL* and *SO×10* were highly upregulated within the tumor programs (Figure 5d). Similar to the pathologist annotations, the model also detected spatial programs corresponding to stromal (fibroblast) and lymphoid tissues (Figure 5c), which marker genes including *COL1A1* (fibroblast-specific, Figure 5c-d) and *CXCL13* (lymphoid-specific, Figure 5d). Notably, *MYL9* was one of the most highly expressed genes within the fibroblast expression signature (Figure 5d), which is a marker of tumor-associated myofibroblasts^35^, indicating that the fibroblast program identified by BayesTME represents a subpopulation of fibroblasts reprogrammed by their proximity to the tumor. In the fibroblast-related spatial program, immune-related hub genes like *IGLL5* and *IGJ* displayed an enrichment at the tumor boundary (Figure 5c). The model also identified a macrophage-related spatial program (Figure 5c), which had not been detected by the pathologist. One of the top macrophage marker genes, *CXCL9* (Figure 5c-d) is a marker of tumor-associated macrophages^36^, which have an important role in anti-tumor immunity^37^. Taken together, our results show that BayesTME can successfully not only recapitulate, but also improve the detection of novel tumor and tumor-associated cell types that are difficult to identify purely by histology.

### BayesTME discovers spatial programs capturing muscular bilateral symmetry and tumor-immune interaction in a zebrafish melanoma model

We expanded upon our human melanoma results by applying BayesTME to our recently published dataset of zebrafish *BRAF*^*V* 600*E*^ -driven melanoma^23^, generated using the 10X Genomics Visium technology with approximate spot resolution of 55 *µm*. Both samples contained tumor and TME tissues (muscle, skin) (Figure 6, Figure 7).

**Figure 6:**
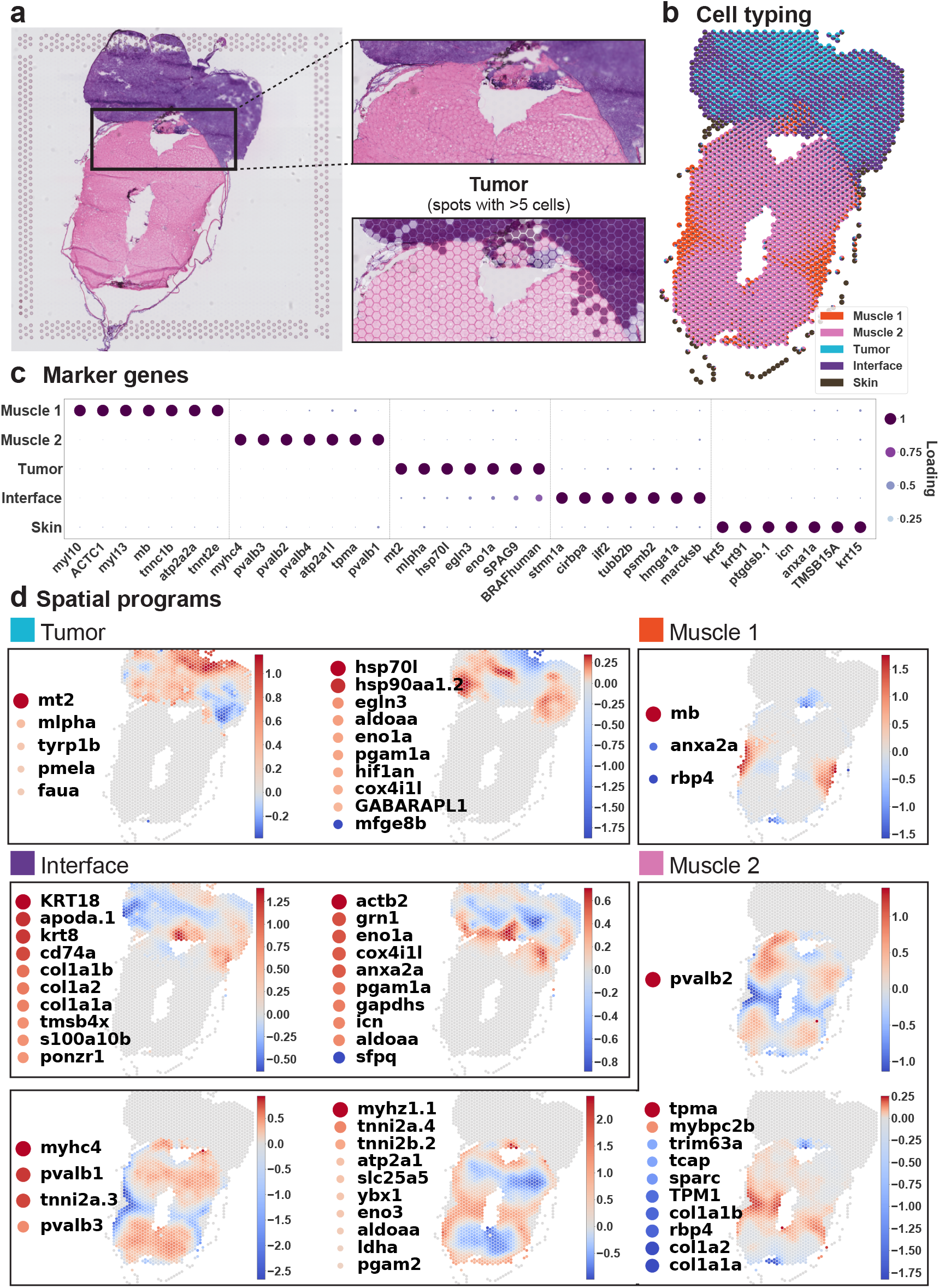
BayesTME identifies sharp boundaries and novel tumor interface programs in a zebrafish melanoma model. (**a**.) Histology of zebrafish sample A; cutout: zoom in on the tumor interface region. (**b-c**.) BayesTME discovers 5 cell phenotypes with biologically plausible marker genes; cutout: zoom in on the recovered tumor/not-tumor proportions show BayesTME captures the sharp tissue change point. (**d**.) 9 spatial transcriptional programs discovered at a 5% FDR; muscle programs illustrate BayesTME captures bilateral symmetry without prior knowledge; interface and tumor programs capture differences between interior and exterior tumor behavior.

**Figure 7:**
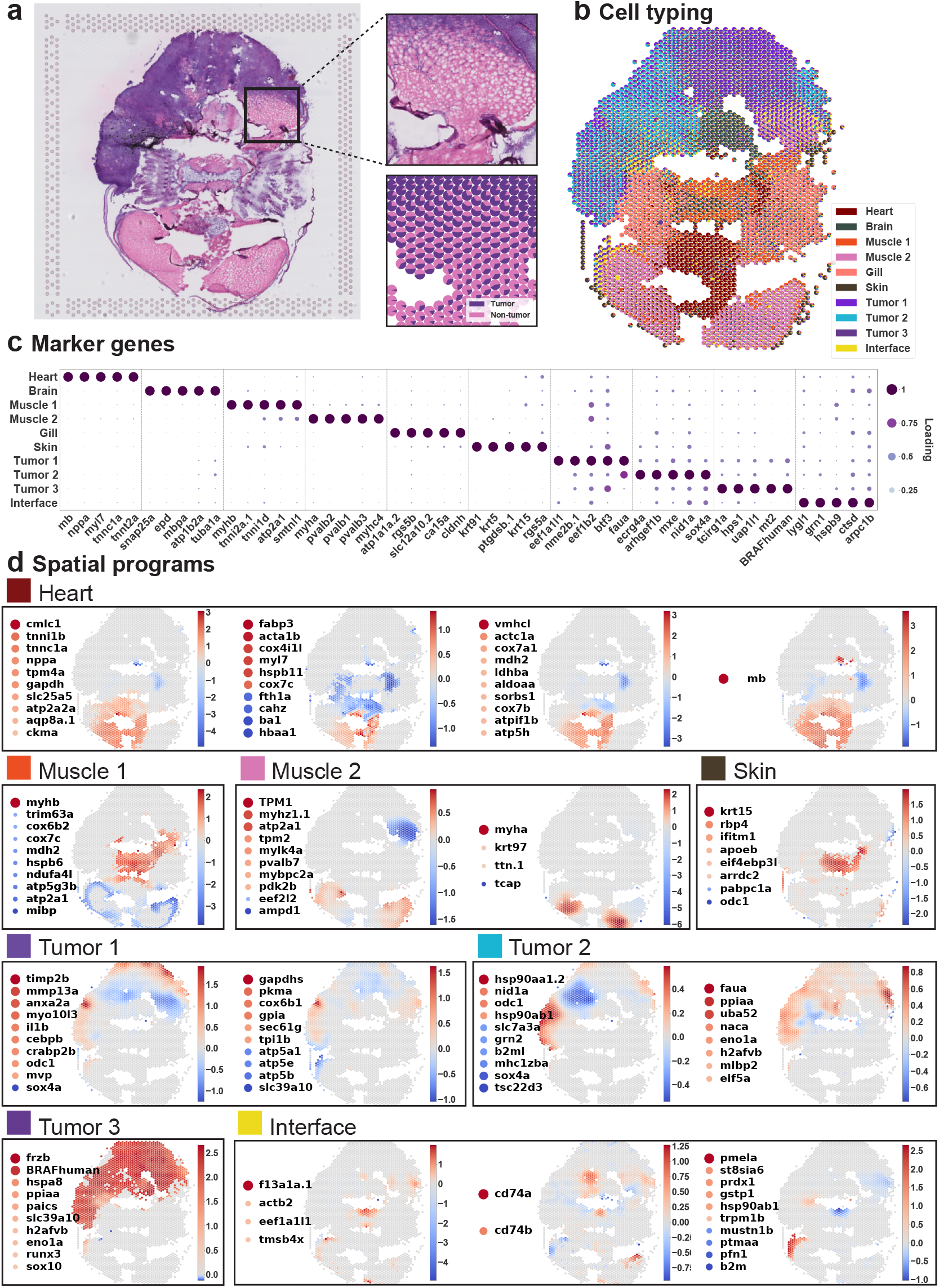
BayesTME reveals gradual tumor invasion and confirms interface programs in a second zebrafish melanoma model. (**a**.) Histology of the zebrafish B sample; cutout: tumor interface with gradual invasion of tumor cells into the muscle region. (**b**.) BayesTME cell types recovered; cutout: corresponding tumor-muscle interface with tumor/non-tumor proportions capturing the gradient of tumor invasion present in histology. (**c**.) Marker genes for the discovered 10 cell types. (**d**.) 16 spatial transcriptional programs discovered at a 5% FDR.

Within Sample A, BayesTME identified cell types corresponding to tumor, skin, and muscle (Figure 6b-c). Each cell type upregulated expected marker genes, such as myosins and parvalbumins in muscle (*myhc4, myl10, pvalb1, pvalb2, pvalb3, pvalb4*), *BRAF*^*V* 600*E*^ in tumor, and keratins in skin (*krt5, krt91, krt15*) (Figure 6c). Two celltypes (“Tumor” and “Interface”) were detected within the tumor, both expressing *BRAF*^*V* 600*E*^ (Figure 6a-c). Although the tumor region of Sample A bordered adjacent muscle with little mixing of the two tissue types visible on the H&E-stained section, the interface cell type appeared to infiltrate into the neighboring TME, reminiscent of the interface cell we identified in our recent work^23^ (Figure 6a). Many of the interface marker genes were the same as interface marker genes we previously identified, including *stmn1a, tubb2b*, and *hmga1a*^23^ (Figure 6c). Both spatial programs corresponding to the interface type were enriched at the tumor boundary (Figure 6d). In addition to the interface marker genes we previously identified, BayesTME uncovered a number of genes related to remodeling of the extracellular matrix (ECM) that displayed a spatial enrichment at the tumor boundary, including several collagen-related genes (col1a1a, col1a2, col1a1b; Figure 6d), consistent with a role for the interface cell state in melanoma invasion. Immune genes were also enriched at the tumor-muscle interface, including *ilf2* and *grn1* (Figure 6c-d).

Sample B contains a wider variety of tissue types including heart, brain, gills, tumor, and muscle (Figure 7a-c). Mixing of tumor and muscle tissues at the tumor boundary was visible by histology (Figure 7a). Notably, BayesTME again uncovered an “interface” cell state specifically enriched at the tumor boundary (Figure 7a-b). Similar to Sample A, a number of immune-related genes were spatially patterned and/or enriched in the interface region, including *lygl1, grn1, cd74a/b*, and *b2m* (Figure 7c-d). Melanoma is a highly immunogenic cancer whose interaction with immune cells in the TME significantly influences tumor progression^38^. Whether the enrichment of immune genes at the tumor-TME interface represents pro-inflammatory tumor cells at the tumor boundary, or a type of novel tumor-associated immune cell type will be an exciting topic of future investigation.

In both samples, we uncovered a significant degree of spatially-patterned tumor heterogeneity. BayesTME identified spatial programs characterized by up-regulation of classical melanoma markers such as *pmela* and *tyrp1b* (Sample A “Tumor”, Figure 6d) and *BRAF*^*V* 600*E*^ and *so×10* (Sample B “Tumor 3”, Figure 7d). Other spatial programs identified in the tumor likely represent other facets of tumor biology. Hypoxia-related genes (*hif1an, egln3* ; Figure 6d) were spatially enriched within the tumor region of Sample A, which may indicate hypoxic regions of the tumor due to lack of oxygen supply. Hypoxia has been linked to melanoma progression^39^. We also identified spatially-patterned signatures of metabolism, which could represent different metabolic pathways active within the tumor. One of the spatial programs identified within the tumor region of Sample B up-regulated several genes corresponding to ATP synthase subunits (*atp5a1, atp5e, atp5b*) and other metabolic genes (*gpia, tpi1b*) (Figure 7d). Determining how different metabolic pathways are spatially-organized and regulated within the tumor will be an interesting area of further study. Taken together, our results indicate that BayesTME identifies complex spatial patterns of transcriptional heterogeneity within melanoma and the melanoma microenvironment, and uncovers a potentially novel pro-inflammatory cell state present at the tumor boundary.

## Discussion

This paper has presented BayesTME, a reference-free Bayesian method for end-to-end analysis of spatial transcriptomics data. Compared with existing scRNA-seq referenced methods, BayesTME applies to a wider variety of tissues for which scRNA-seq may not be tractable due to economic, technical, or biological limitations. Even when references are available, highly heterogeneous and diseased tissues may contain different subsets of cell types between consecutive samples. However, BayesTME is adaptable to scRNA-seq reference if a reliable one is available. With reference data, one can obtain the empirical estimation of the expression signature *ϕ*, which is invariant to sequencing depth batch effects. Computationally, access to pre-clustered scRNA-seq significantly accelerates the inference by removing the need to perform cross-validation to select the cell phenotypes. On the other hand, unlike most reference-free methods, BayesTME does not rely on dimension reduction like PCA. This advantage enables BayesTME to draw individual gene-level inferences including expression signatures, phenotype markers, and spatial transcriptional programs which current methods miss. Our comparison to 11 other ST data analysis methods highlighted BayesTME’s advance in bleed correction, spot deconvolution, tissue segmentation, and within-cell-type spatial spatial variation in gene expression.

Advances in ST technology promise to soon enhance the resolution to near-single cell levels, dramatically increasing the number of spots. We have carefully designed the computational inference routines in BayesTME to meet this challenge. BayesTME scales sub-linearly with the number of spots, with a 100x increase in the number of spots leading to only a 10x increase in computational runtime (Supplementary Fig. 9). To further speed up inference, one can place an informative prior on the cell count in a given spot using the H&E slide as reference; simulation experiments show that with a handful of noisy cell count annotations, the cell count accuracy also improves to nearly perfect (see the Supplement for details).

Understanding how cells alter their expression levels as a function of their spatial location in a tissue is necessary for a complete characterization of the cellular architecture of the tissue microenvironment. BayesTME captures these expression level changes in the form of spatial transcriptional programs. Our results showed BayesTME is able to capture biologically meaningful spatial programs which hint at cell-cell interaction in tumor microenvironments. To further facilitate our understanding of cell-cell interaction mechanisms, future versions of BayesTME will introduce an additional cell-type interaction term in the success rate formulation in our negative binomial model. This interaction term will model the total influence cell type *k* in spot *i* as the sum of the interactions between cell type *k* and all possible cell types *k*^′^. We also plan to explore extending this formulation to all spots within a reasonable neighboring of spot *i* for global interactions triggered by paracrine, synaptic, or endocrine signaling. This process is computationally expensive under the current ST technology. However, with single-cell resolution, such inference becomes tractable as we only need to look at the individual cells of different cell types within the reasonable neighborhood of cell Increased ST resolution will significantly drop the computation cost by a factor of *K*, which can also be vectorized to further speed up this process. Thus, BayesTME is well-positioned to make future computational advances in ST modeling, in step with the coming technological advances in ST methods.

## Methods

### Notation and setup

We assume we are given an *N* × *G* matrix *R* where *R*_*ig*_ is the UMI counts for gene *g* at spot *i*. The spot *I* is associated with some known location *l*(*i*) ∈ ℝ^2^ on the tissue. These locations define a graph 𝒢 = (𝒱, ℰ) where each vertex is a spot. There is an edge between two vertices if they are within some *ϵ* distance. We set 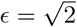 such that each non-boundary spot has 4 neighbors for lattice layouts (e.g., Slide-seq) and 6 neighbors for hexagonal layouts (e.g., Visium). We assume that there are *K* cell phenotypes (hereon simply called cell types) in the sample, each with their own expression profile. We do not assume that *K* is known nor do we assume that there is side information about different cell types and their expression profiles (i.e., we do not assume access to paired single-cell RNA). We refer to UMI counts and read counts interchangeably, where read counts are understood to mean UMI-filtered reads and not raw, possibly-duplicated reads.

### Generative model

BayesTME models several sources of spatial variation in ST data using a single hierarchical probabilistic model,

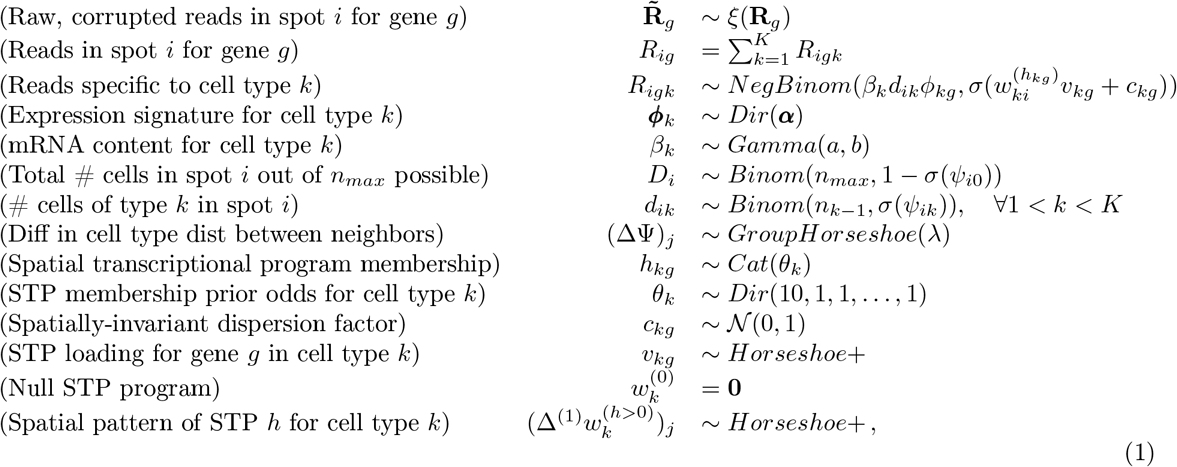

where *σ* is the logistic function, *D*_*i*_ is the total number of cells in spot *i*, and *λ* is the hyperparameter that controls the degree of spatial smoothing. The function *ξ*(·) is a nonparametric function defining the spot bleeding process that probabilistically maps from the true read counts **R**_*g*_ for each gene *g* to the observed counts 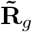. We specify no functional form for this function and only constrain it to be decreasing in the distance from the true to observed spot location. The matrix ∆ is the edge-oriented adjacency matrix encoding the spot graph, also equivalent to the root of the graph Laplacian; ∆^(1)^ = ∆^*T*^ ∆ is the first-order graph trend filtering matrix^40,41,^ equivalent to the graph Laplacian.

Since full Bayesian inference in the above model is computationally intractable, we develop an efficient empirical Bayes approach that splits posterior inference into stages. This piecewise approach to fitting is distinguished from the ad hoc pipeline approach of existing workflows in that a single, coherent generative model is driving the estimation. The empirical Bayes approach merely plugs in point estimates for nuisance parameters while providing full Bayesian inference with uncertainty quantification for the latent variables of interest.

### Gene selection

BayesTME scales linearly with the size of the gene library. To keep posterior inference computationally tractable, we select the top *G* = 2000 genes ordered by spatial variation in log space. Specifically, we transform the reads as log(1+*R*) and rank each column by the variance, keeping the top 2000. The logarithmic transform separates spatial variation from natural variation that arises due to simply having a higher overall expression rate. We then drop all ribosomal genes (i.e., those matching an ‘rp’ regular expression). After selecting and filtering the top genes, we work directly with the UMI read counts.

### Anisotropic bleed correction

Technical error causes UMIs to bleed out from barcoded spots. BayesTME models this bleed as a combination of unknown global and local effects. Global effects form a baseline bleed count for any spot, corresponding to a homogeneous diffusion process. Local effects imply that the UMI count at a given spot is a function of how far it is from the original location of each of the UMIs. BayesTME employs a semi-parametric, anisotropic model for global and local effects,

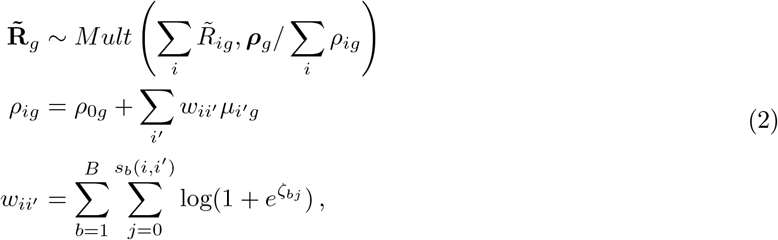

where 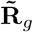 are the raw, observed counts and *ρ*_0*g*_ are the global effects. The local effects in Equation (2) are modeled using a set of *B* monotone nonparametric basis functions *ζ* that decay as a function of the basis-specific pseudo-distance *s*_*b*_.^2^ BayesTME uses the four cardinal directions (North, South, East, and West) for the basis functions. This choice is based on the observation that UMIs tend to bleed toward one corner. We also observed that bleeding appears to be less extreme in tissue regions than non-tissue regions. Thus, BayesTME distinguishes between in- and out-of-tissue distance by learning four separate basis functions for each region. The distance from an original spot *i*′ to its observed spot *i* is then a summation of the in- and out-of-tissue components of a straight line between the two spots.

The bleeding model is fit by alternating minimization. At each iteration, BayesTME alternates between estimating the basis functions 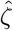 and global rates 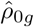, and estimating the latent true UMI rates 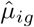. After the model is fit, BayesTME replaces the raw reads with the approximate maximum likelihood estimate of read counts,

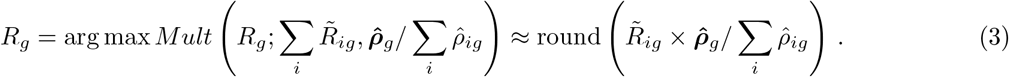

The cleaned reads *R* are then treated as correct in subsequent inference steps. This can be seen as an empirical Bayes approach, where the model in Equation (2) is optimized and uncertainty over *R* is replaced with a point estimate that maximizes the marginal likelihood of possible true read configurations.

### Discrete deconvolution model

The spot-wise gene counts *R*_*ig*_ can be decomposed into the sum of cell type-specific gene reads in any given spot, i.e.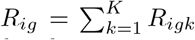. BayesTME models the cell type-specific reads with a Poisson distribution controlled by three parameters *β*_*k*_, *d*_*ik*_ and *ϕ*_*kg*_. Specifically, *β*_*k*_ denotes the expected total UMI count of individual cells of type *k*; *d*_*ik*_ denotes the number of cells of type *k* located in spot *i*; and ***ϕ***_*k*_ = (*ϕ*_*k*1_, …, *ϕ*_*kG*_) denotes the gene expression profile of cell type *k*, where each element *ϕ*_*kg*_ is the normalized expression of gene *g* in cell type *k*; equivalently, *ϕ*_*kg*_ is the proportion of UMIs that cell type *k* allocates to gene *g*. The generative model for BayesTME follows,

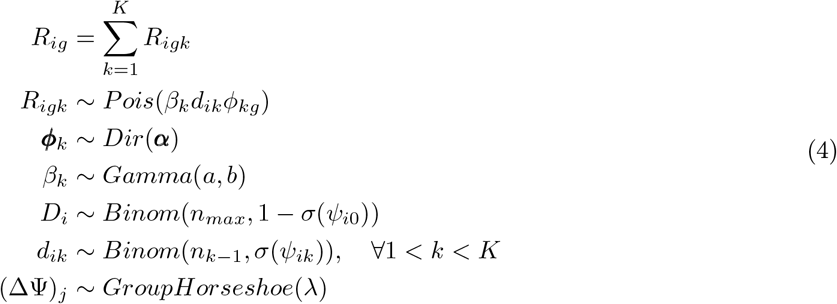

where *D*_*i*_ is the total number of cells in spot *i*, and *λ* is the hyperparameter that controls the degree of spatial smoothing. The matrix ∆ is the edge-oriented adjacency matrix encoding the spot graph 𝒡, also equivalent to the square root of the graph Laplacian. The hierarchical prior encoded by the last three lines of Equation (4) is a heavy-tailed Bayesian variant of the graph-fused group lasso prior^27,26^ that uses the Horseshoe+ distribution^25^. This prior encourages the probability distribution over cell type proportions to follow a piecewise constant spatial function, encoding the prior belief that cells form spatially contiguous communities. The model is data-adaptive, however, and able to handle deviations from this prior where warranted in the data; see for example, the smooth gradient of cell type proportions recovered in Figure 7.

### Posterior inference

Posterior inference in BayesTME is performed through Gibbs sampling. The full derivations for all complete conditionals and update steps are available in the supplementary material. The key computational innovations in BayesTME come in the form of a fast approach to update *d*_*ik*_, the number of cells of type *k* in spot *i*. As we show in the supplement, block joint sampling over all **d**_*i*_ and *D*_*i*_ can be done via an efficient forward-backward algorithm. This algorithm effectively converts the cell count prior to a hidden Markov model prior. The Poisson likelihood in Equation (4) acts as the emissions step and the emission log-likelihood can be collapsed into a series of fast updates. This inference step enables us to sample over the entire combinatorial space of possible cell counts in 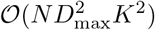 time for *N* spots, *K* cell types, and 0 ≤ *D*_*i*_ ≤ *D*_max_ possible total cells in each spot. BayesTME performs Gibbs sampling using these fast updates with a burn-in and Markov chain thinning; we use 2000 burn-in steps, 5 thinning steps between each sample, and gather a total of *T* = 100 post-burn-in posterior samples.

### Selecting the number of cell types and smoothness hyperparameters

BayesTME automatically chooses the number of cell types *K* via *M* -fold cross-validation. For each fold, a random non-overlapping subset of the spots are held out; we use *M* = 5 folds with 5% of spots held out in each fold. The spatial priors in BayesTME enable imputation of the cell type probabilities at each held out spot in the training data. For each fold, we fit over a discrete grid of *λ* smoothness values; we use *λ* = (10^0^, 10^1^, …, 10^6^). For a given fold *m*, cell type count *K*, and smoothness level *λ*, we calculate the approximate marginal log-likelihood of the held out spots using *T* posterior samples,

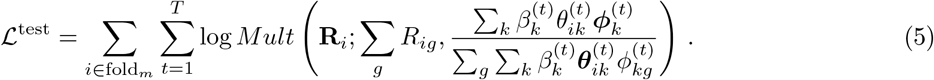

Results are averaged over all *λ* values for each fold and then averaged across each fold. The *λ* averaging is an empirical Bayes estimate with a discrete prior on *λ* integrated out; the cross-validation averaging is an unbiased approach to selecting *K*. After selecting *K*, we refit BayesTME on the entire data using the chosen *K* and the *λ* with average cross-validation log-likelihood closest to the overall average.

### Selecting marker genes

We define a gene as a marker of a particular cell type if its expression in that cell type is significantly higher than in any other cell type. BayesTME uses posterior uncertainty to select statistically significant marker genes with control of the Bayesian false discovery rate (FDR)^42^. To calculate the local FDR we use the *T* posterior samples,

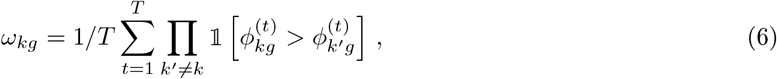

yielding the posterior probability that gene *g* is a marker for cell type *k*. We sort the *ω* values in descending order and solve a step-down optimization problem,

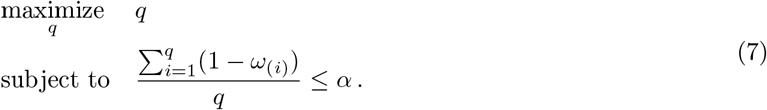

The set of *ω* values selected controls the Bayesian FDR at the *α* level. BayesTME can alternatively control the Bayesian Type I error rate at the *α* level by only selecting marker genes satisfying *ω*_*kg*_ *≥* 1 − *α*. We then rank the selected marker gene candidates by *ω* and *ξ* jointly, where

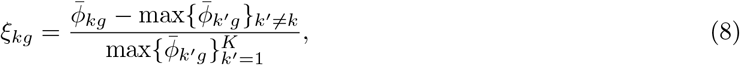

is the normalized expression score in [−1, 1] measuring the expression level of gene *g* in cell type *k* compared with all the other cell types, and 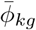 is the posterior mean of *T* posterior samples. By default, we set the FDR threshold to 5%; our results report an interpretable subset of the top 20 genes for each inferred cell type.

### Community detection

To segment the tissue into cellular communities, BayesTME clusters the fused spatial probabilities **Ψ**. First, the neighbor graph is augmented with the nearest 10 neighbors to adjust for spatially-disconnected spots due to tissue tears in sectioning. The posterior samples are flattened into a single vector for each spot. Spots are then clustered using agglomerative clustering with Ward linkage, as implemented in `scikit-learn`. The number of clusters *q* is chosen over a grid of *q ∈*(1, …, 50) to minimize the sum of the AIC^43^ and BIC^44^ scores. Community distributions are calculated as the average of all posterior probabilities of all spots assigned to the community. When comparing community segmentation in benchmarks, we applied BayesTME’s clustering algorithm on DestVI and stDeconvolve, as they do not provide segmentation routines.

### Spatial transcriptional program model

The deconvolution model in Equation (4) assumes gene expression is stationary within a given cell type. However, we expect that variation in a small number of important genes should be spatially dependent. BayesTME captures this spatial variation by replacing the Poisson likelihood in Equation (4) with a more complex negative binomial one,

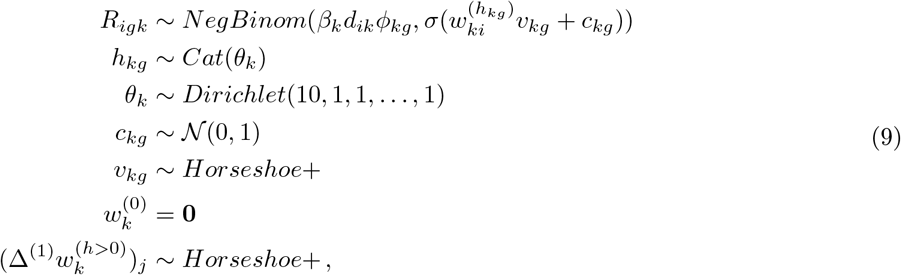

where *σ* is the logistic function and ∆^(1)^ = ∆^*T*^ ∆ is the first-order graph trend filtering matrix, equivalent to the graph Laplacian. The rate in Equation (9) is equivalent to that in the simpler model in Equation (4). In both cases, the expected read count scales additively with the number of cells, a crucial property that reflects the intuition that a spot with twice as many cells should yield twice as many reads.

Gene expression within a cell type varies spatially through the success probability (the second term) in the negative binomial likelihood. The offset term *c*_*kg*_ corresponds to the spatially-invariant expression term that controls the dispersion rate in the counts. Each gene *g* in each cell type *k* belongs to one of *H* clusters. Each cluster defines a different spatial pattern 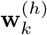, which we refer to as *spatial transcriptional programs*. The first program 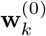 is the null program corresponding to spatially-invariant expression. All subsequent programs are latent and inferred through posterior inference. BayesTME places a heavy prior on genes coming from the null, such that it takes substantial evidence to conclude that a gene is spatially varying within a cell type; this prior is necessary as otherwise the model is only weakly identifiable. Genes that participate in the non-null spatial programs do so by placing a weight *v*_*kg*_ on the spatial pattern. This weight shrinks, magnifies, or can even invert the pattern, allowing for clustering of negatively correlated genes into the same spatial transcriptional program. BayesTME places a sparsity-inducing prior on *v*_*kg*_ in order to encourage only strongly-participating genes to be assigned to non-null programs.

### Spatial transcriptional program inference

Posterior inference via Gibbs sampling is possible with the STP BayesTME model. However, the fast HMM updates for the cell counts are no longer available, making the inference algorithm substantially slower. For computational efficiency, we instead take a two-stage approach. First, we fit the deconvolution model in Equation (4), collecting *T* posterior samples of each latent variable. Then we fix (*β*, **d**, Φ)^(*t*)^ for each sample *t* = 1, …, *T*. For each fixed sample, we run a new Gibbs sampler for the non-fixed variables in Equation (9); we use 99 burn-in iterations and take the 100^th^ iteration as the sample for the *t*^th^ iteration of the full model parameters. We motivate this approach mathematically by the identity that if *Y ∼ Pois*(*r*) and *X ∼ NegBinom*(*r, p*) then 𝔼[*Y*] = 𝔼 [*X p* = 0.5]. Since we put sparsity priors on *v*_*kg*_ and a standard normal prior on *c*_*kg*_, all of our priors are peaked at *p* = 0.5. Thus, *a priori*, we expect the posterior mean under the full joint inference model to be nearly the same as the two-stage model; in practice, we find the two approaches produce similar results.

### Selecting significant spatial transcriptional programs

Spatial transcription programs in BayesTME correspond to spatial patterns in 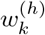 in cell type *k* and the members of a spatial program are the genes *g* for which *h*_*kg*_ is significantly non-null. Spatial programs are only considered active in spots *i* where *d*_*ik*_ *>* 0 with high probability. Specifically, for a given *α* significance level, we select spots and genes for spatial program *s* in cell type *k* as follows,

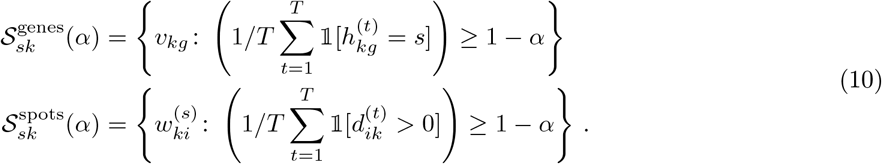

If either 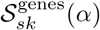 or 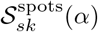 is empty, we filter out the entire program. We also filter any programs where the Pearson correlation between 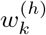 and *d*_*k*_ is more than 0.5 and Moran’s *I* spatial autocorrelation less than 0.9; these programs capture technical noise and overdispersion rather than meaningful spatial signal. In practice, we find *H* = 10 to be a sufficient number of potential spatial programs per cell type. BayesTME sets the spatial transciptional program significant threshold to *α* = 0.95.

## Footnotes

1 https://github.com/tansey-lab/bayestme

2 Technically these basis functions are pseudo-distances as they do not satisfy symmetry and thus are not metric functions.

