## Supplement for "BayesTME: A unified statistical framework for spatial transcriptomics"

### BayesTME: Supplementary information

#### 1 Deconvolution benchmark simulations

We took the mouse brain scRNA-seq data from cell2location as the reference and picked $K$ cell types at random. We used the spatial layout of the human melanoma sample as the template and partitioned the tissue template into 14 blocks. We drew the cell type proportions for each block from a Dirichlet distribution with concentration parameter $\alpha$ equal to $1/K$. This concentration induces sparsity in the subset of cell types in each spot, which replicates real tissues where cell types are often present only in a subset of the tissue. We sampled the blockwise mean total cell count per spot $\lambda_i$ from $Pois(30)$, and drew the total cell number for each spot from $Pois(\lambda_i)$. With the total cell count and the cell type proportion, we sampled the cell count for each cell type from a multinomial distribution. Given the cell count for each cell type, we sampled the cells from the scRNA-seq data and filled the spots, which results in the semi-synthetic data. We show BayesTME gives accurate inference on cell type probability, cell number, and expression signature (Figure 1 a) and visualization of BayesTME's cell type deconvolution and cellular community segmentation results (Figure 1 b-c) in comparison with the benchmark methods.

#### 2 Spatial transcriptional program benchmark simulations

For the spatial differential expression experiments, we first generated the non-spatially-varying-expression data as above, but using the zebrafish A layout as it has an order of magnitude more spots. We then generated 6 hand-designed spatial patterns. For each cell type, we randomly assigned 2 spatial patterns and randomly selected 5 genes for each spatial pattern. We ranked the selected genes and permuted these genes by matching the ranking to the signal strength in the designed spatial patterns. This approach ensures that each gene has the same marginal distribution as the scRNA-seq clusters. Thus, a spatially invariant cell type inference would detect the same spatial signature as in the non-varying data. BayesTME detects STPs with high accuracy, outperforms the differential expression results from DestVI (Figure 2).

#### 3 Integrating human-annotated H&E cell counts

We conducted a preliminary analysis on the benefits of including a handful of human-annotated approximate cell counts for spots, using the H&E. Such annotations are inherently noisy as the H&E does not perfectly align with the spot locations, leading to potential differences between the H&E location of a spot predicted by the ST platform software and the true location sequenced. To simulate human H&E annotations, we generated a semi-synthetic dataset using the procedure described in Section 1 with 3 cell types. To investigate the help of annotations in cell count inference, we added noisy cell count annotations at $n = 1, 3, 10$ randomly selected spots and compared the performance with the base model without any cell count annotation (Figure 3). Simulating the noisiness of either manual counting or some machine learning algorithm prediction, we made each annotation a random draw from $Pois(D_i^*)$, where $D_i^*$ denotes the ground truth cell count in the chosen spot $i$. With only 1 noisy annotation, BayesTME is able to obtain a significant improvement in cell count accuracy, while the cell count accuracy also improves to nearly perfect with only 10 annotations.

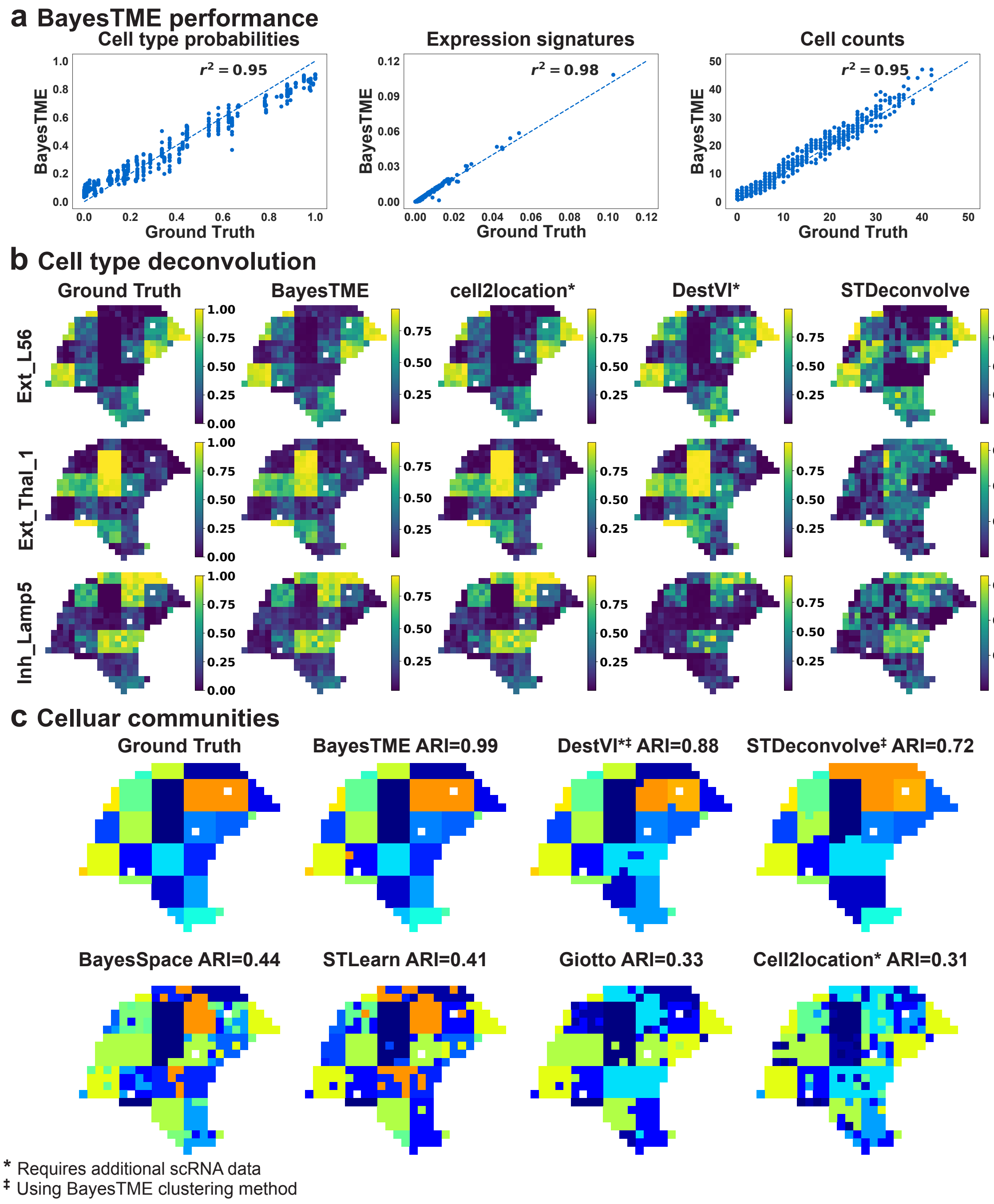


**Figure 1: BayesTME performs favorably to existing methods in semi-synthetic benchmarks. a.** BayesTME accurately recovers the ground truth cell type distributions ($r^2 = 0.95$), discrete cell counts ($r^2 = 0.95$), and expression signatures ($r^2 = 0.98$) of each cell type in the simulation. **b.** Visualization of cell type deconvolution results of BayesTME, compared with reference-based methods cell2location, DestVI, and reference-free methods STdeconvolve. **c.** BayesTME outperforms all existing methods in celluar community segmentation.

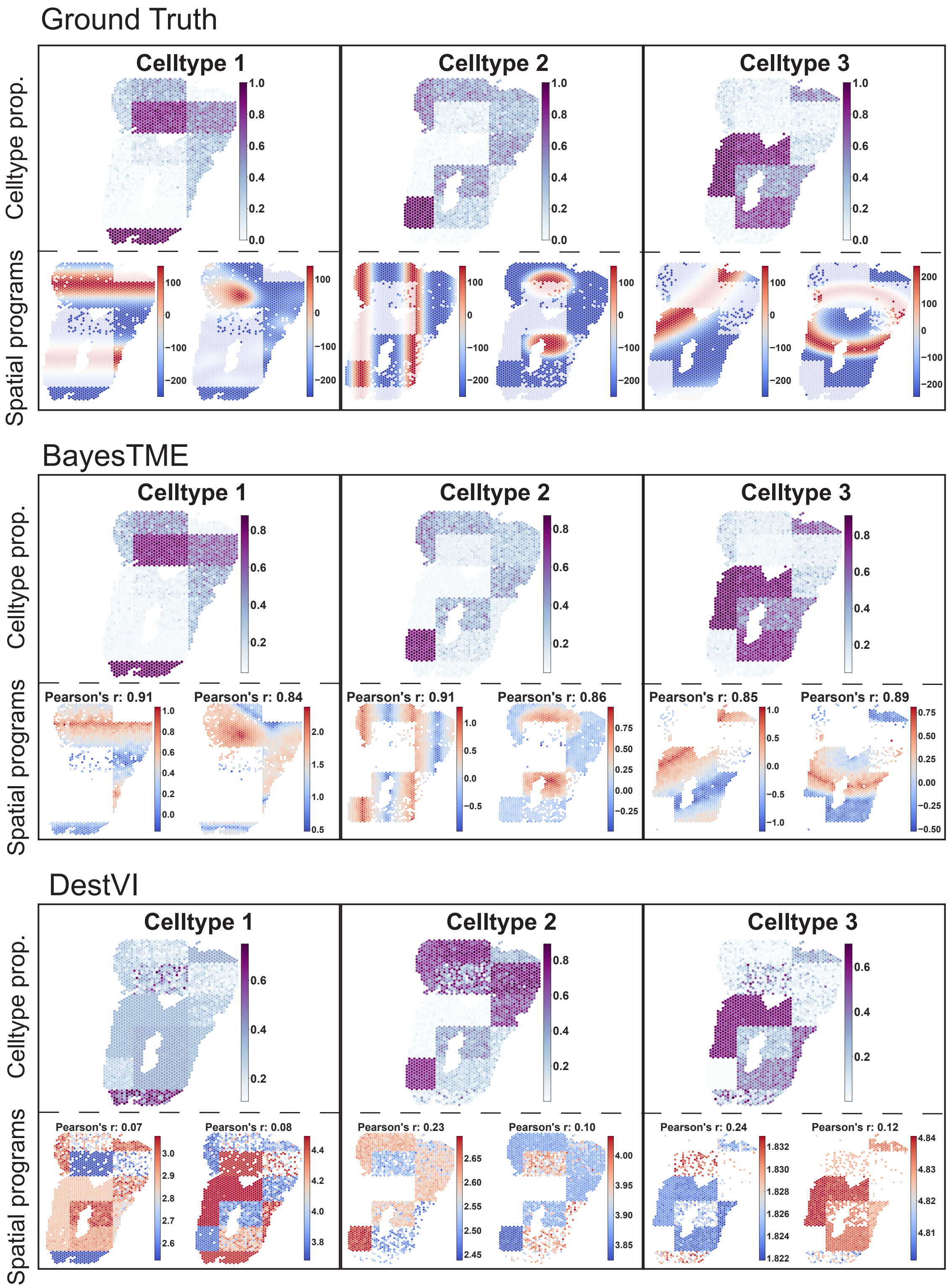


**Figure 2:** Comparison of cell type deconvolution and spatial transcriptional programs detection accuracy between BayesTME and DestVI in simulation dataset with spatial transcriptional programs. BayesTME recovers the ground truth cell number proportion and identifies within-phenotype spatial transcriptional programs with high correlation. DestVI found spatial expression detection results to be uncorrelated with the ground truth.

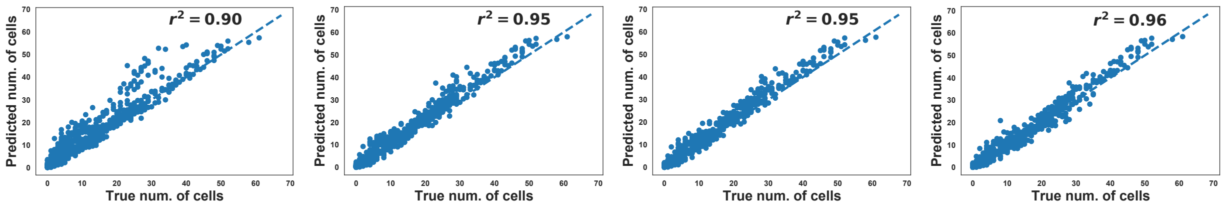


**(a)** 0 annotations **(b)** 1 annotation **(c)** 3 annotations **(d)** 10 annotations

**Figure 3:** Accuracy of discrete cell count inference with 0, 1, 3, and 10 noisy cell count annotations. The performance of the model quickly improves to nearly perfectly capturing the true number of cells.

### 4 Cross-validation selects the correct number of cell types

We constructed 4 semi-synthetic ST data with the same tissue template but different number of cell types ($K^* = 3, 4, 6, 8$). For each semi-synthetic data, we ran cross-validation with number of cell types from 2 to 15 and picked the one with maximum likelihood as our predicted cell type number. The predicted cell type number matches the ground truth $K^*$ in all trials. Such Observation suggest, with cross-validation, BayesTME is able to select the optimal number of cell types from the ST data without scRNA-seq reference.

Figure 4 shows the average cross-validation log-likelihood for each of the four simulations. Each simulation used a different true number of cell types and BayesTME correctly identified the true number for each simulation. As the number of cell types increased, variance in the held out likelihood also increased. Thus, if the number selected is beyond $\approx$ 10 cell types, we recommend increasing the number of folds $m$ to compensate.

### 5 Bleed correction benchmark simulations

The benchmark bleeding simulations use a $70 \times 70$ simulated ST tissue block, roughly the size of a common real ST tissue. We ensure that the tissue region has a 10-spot margin on all sides, capturing the idea that tissue segments should be inscribed well inside of the fiducial markers. To further mimic real tissue, we randomly insert a tissue gap inside the tissue region, since most tissues are not perfectly contiguous.

We set the baseline expression of each gene to be a constant drawn from a gamma distribution with shape 2 and rate 100. We then sample reads for each in-tissue spot from a Poisson distribution with the corresponding rate. We then apply a stochastic bleeding process to corrupt the true UMIs.

We benchmark the decontaminating ability of BayesTME and SpotClean under three different bleeding patterns:

1. **Gaussian bleeding.** Original UMI locations are corrupted by adding noise drawn from a 2d Gaussian with covariance matrix
$$\Sigma_{\text{Gauss}} = \begin{bmatrix} 5 & 1 \\ 1 & 5 \end{bmatrix}.$$
This induces thin-tailed, symmetric bleeding.

2. **Student's-t bleeding.** Original UMI locations are corrupted by adding noise drawn from a 2d Student's-t distribution with scale matrix,
$$\Sigma_{\text{Gosset}} = \begin{bmatrix} 20 & 3 \\ 3 & 30 \end{bmatrix}$$
and 10 degrees of freedom. This induces heavy-tailed, symmetric bleeding.

3. **Anisotropic bleeding.** Original UMI locations are corrupted by adding anisotropic noise that mimics bleeding seen in real tissues. A force 2d vector of $(105, 52.5)$ (150% and 75% of the dimensions of the

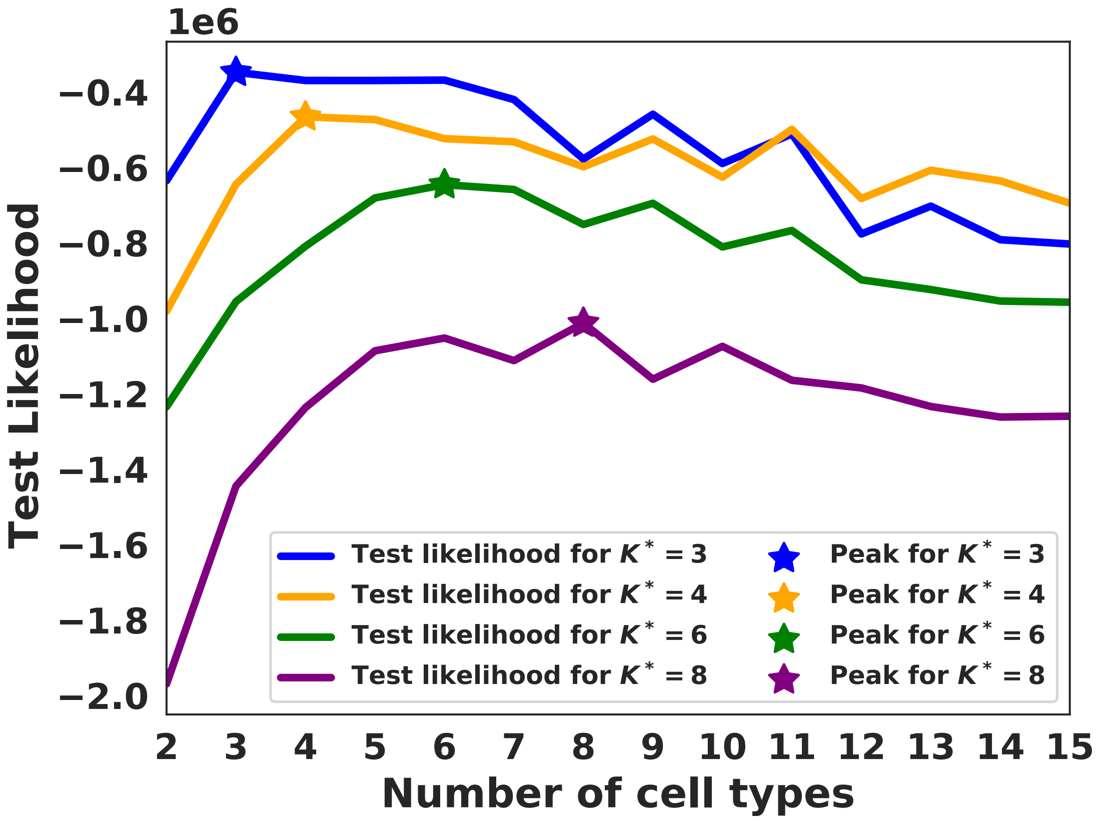


**Figure 4:** The average cross-validation log-likelihood for true number of cell types $K^* = 3, 4, 6, 8$

slide, respectively) is added to each UMI location. A tissue friction coefficient of 5 is used to slow down bleeding within the tissue. Bleeding likelihood then is proportional to this skewed distance via a Laplace kernel with bandwidth 40. See the supplemental code function `distance_weights` in the bleeding simulation script for exact computational details. This creates heavy-tailed, asymmetric bleeding that more closely resembles bleeding seen in ST data than the other two methods.

### 6 Inferred bleeding patterns from zebrafish samples

Following the BayesTME pipeline, we applied the BayesTME preprocessing pipeline to both zebrafish melanoma model samples. Figures 5 and 6 show the inferred directional bias of the bleeding inferred by BayesTME. The results align with the visual observation that in both samples UMIs tend to bleed to a specific direction and bleeding is less substantial inside the tissue than outside. For further insight into the debleeding results, Figures 7 and 8 show the inferred most likely direction of the uncorrupted spot for a UMI observed every spot in the samples.

### 7 BayesTME scales efficiently to ultra-high resolution ST data.

Next-generation ST technologies promise to deliver $2\mu m$ resolution, resulting in an increase of up to two orders of magnitude over the current number of spots. We have carefully designed BayesTME so that it scales to meet these new challenges. Specifically, the main computational step in posterior inference in BayesTME

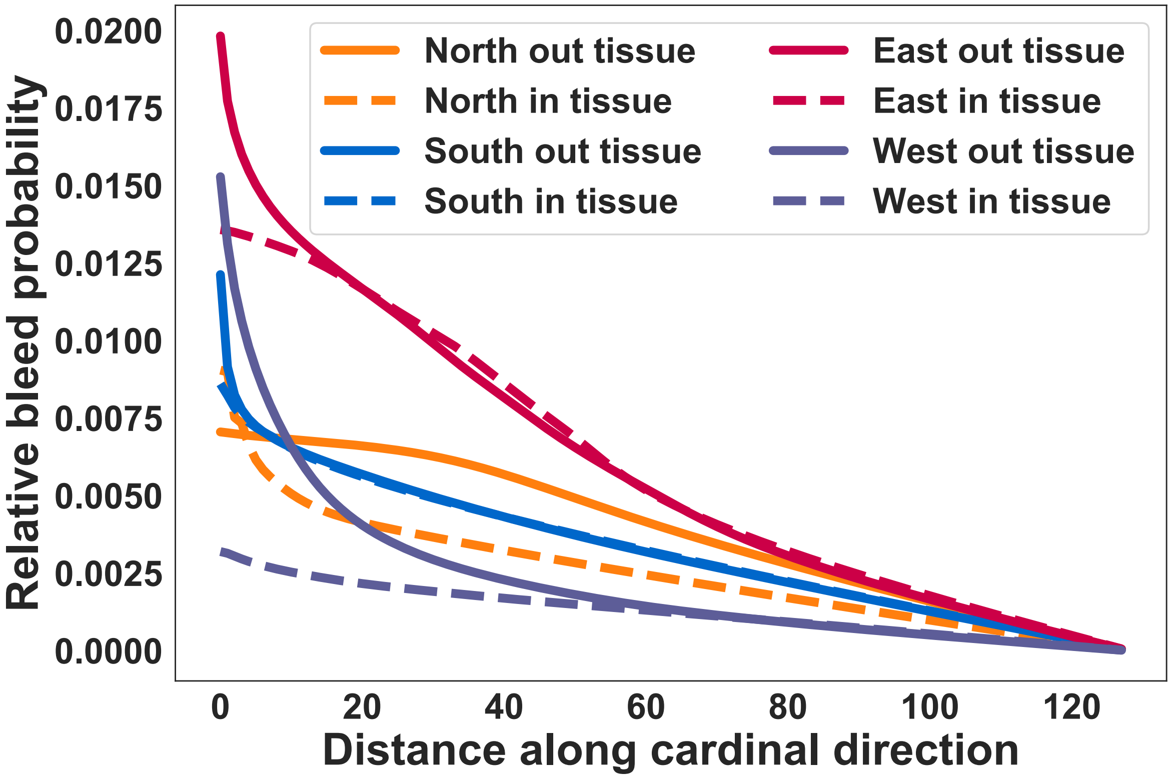


**Figure 5:** Inferred bleeding basis functions for each of the four cardinal directions both inside and outside of the tissue for zebrafish A.

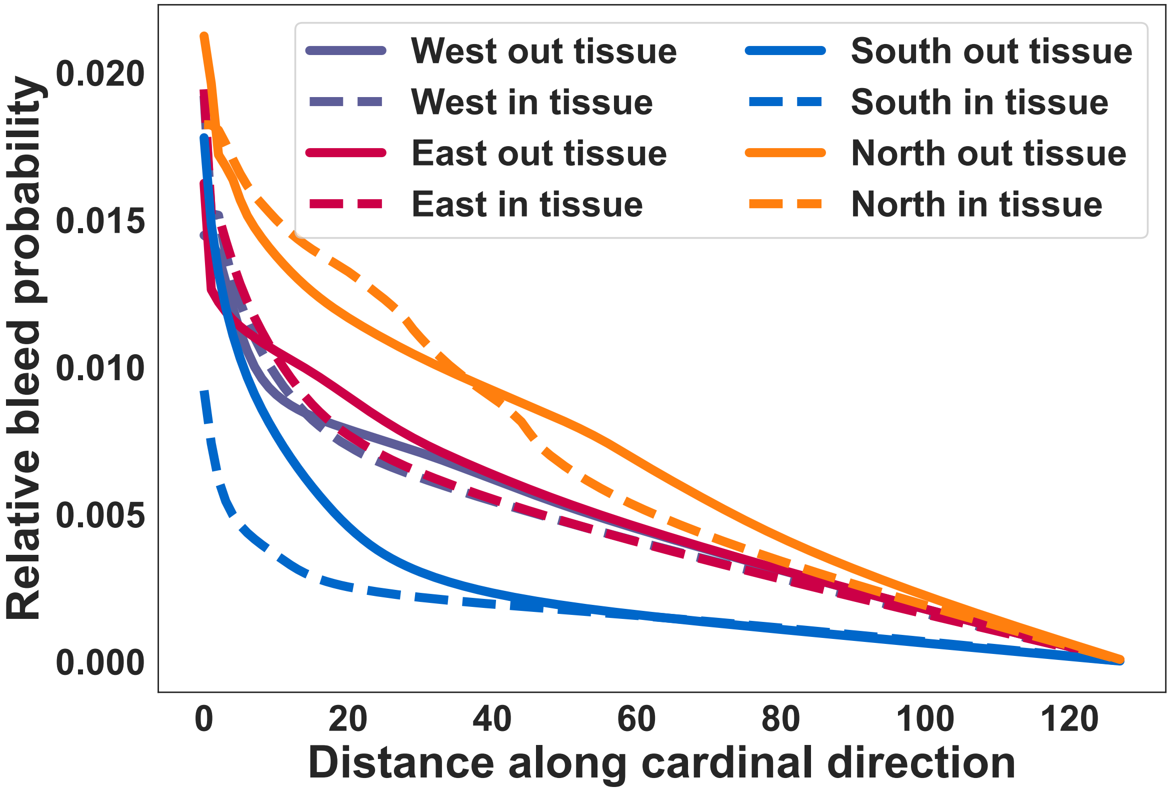


**Figure 6:** Inferred bleeding basis functions for each of the four cardinal directions both inside and outside of the tissue for zebrafish B.

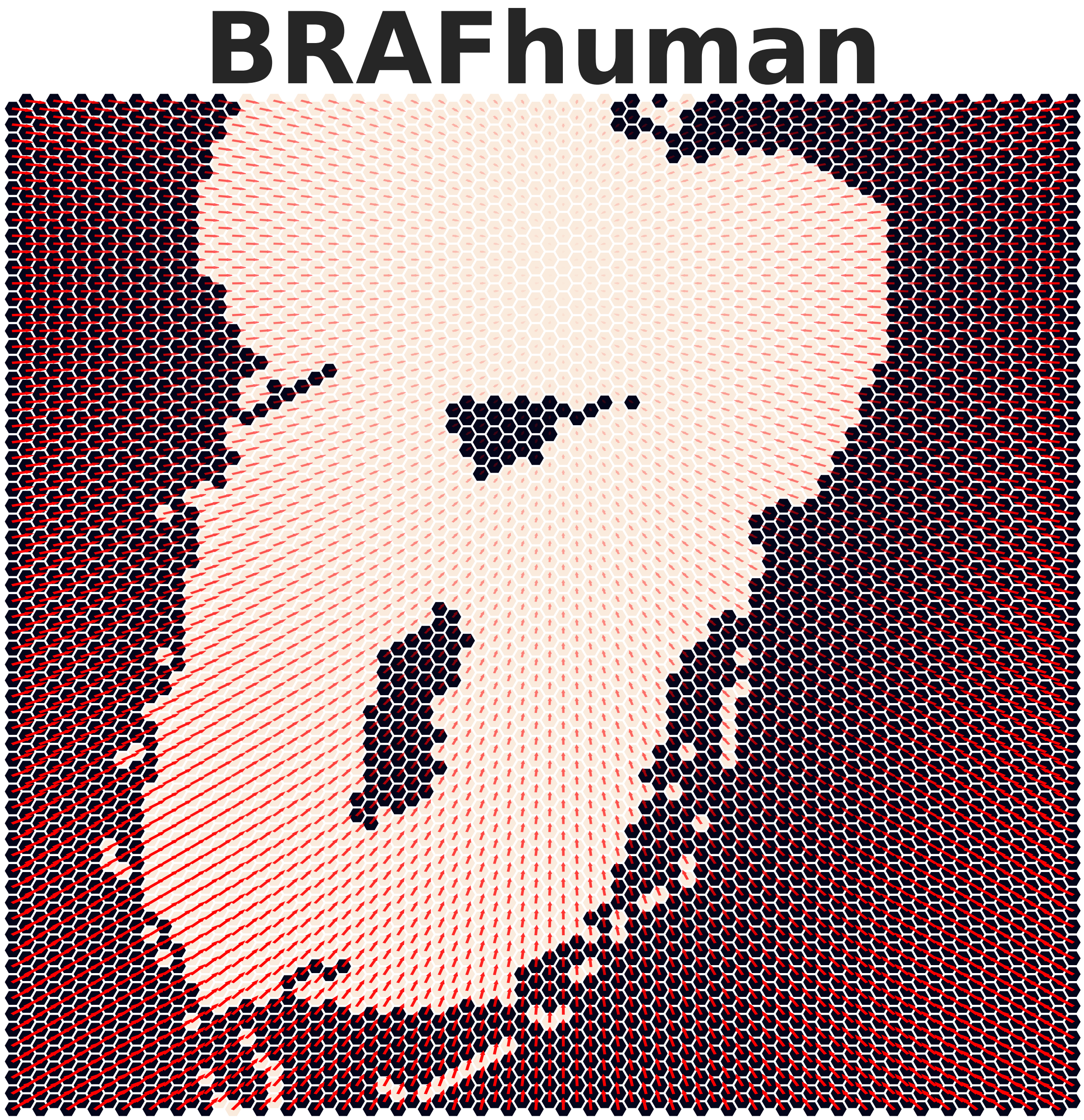


**Figure 7:** Patterns mapping where a UMI of $BRAF^{V600E}$ most likely came from in the zebrafish A sample.

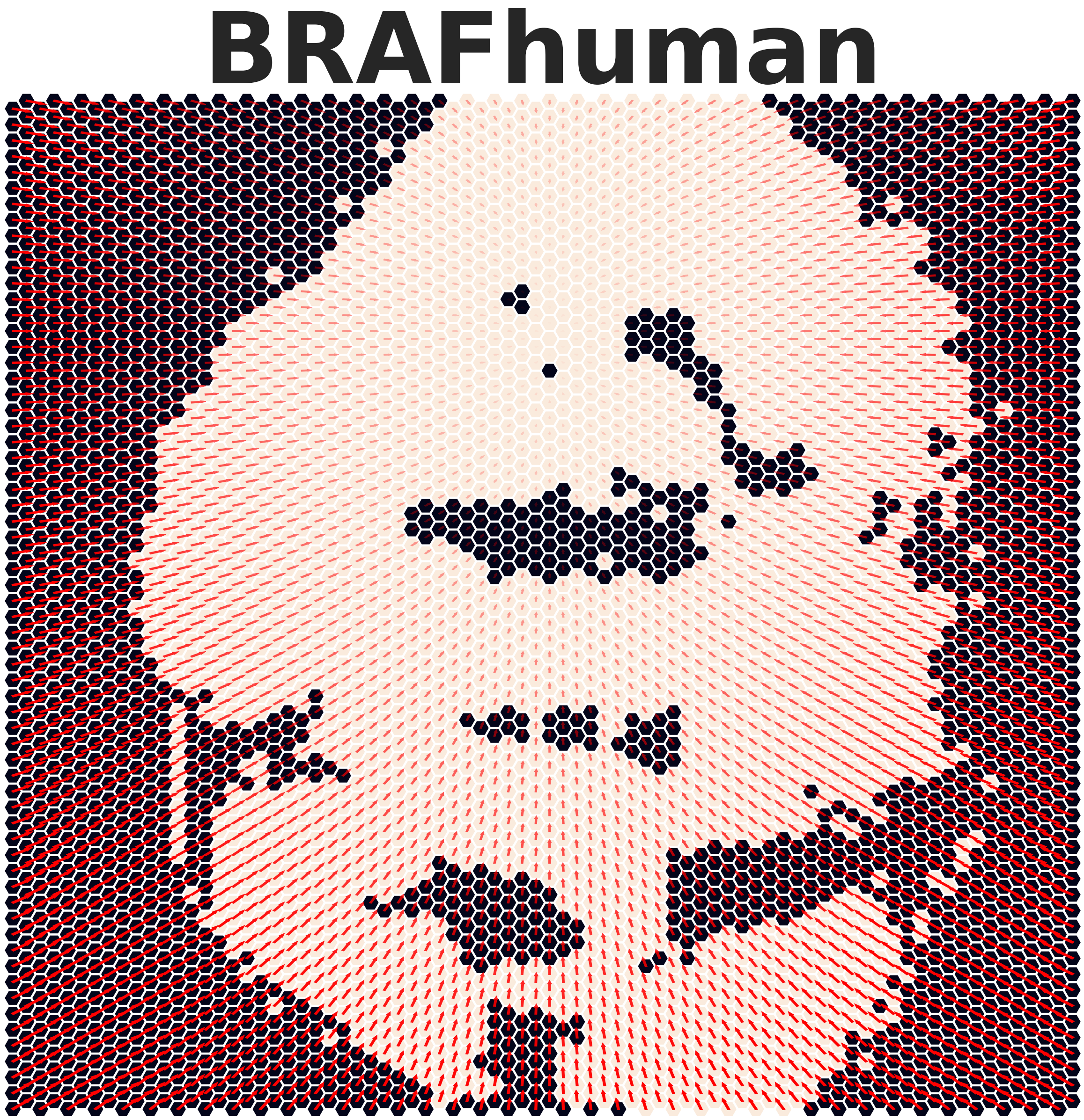


**Figure 8:** Patterns mapping where a UMI of $BRAF^{V600E}$ most likely came from in the zebrafish B sample.

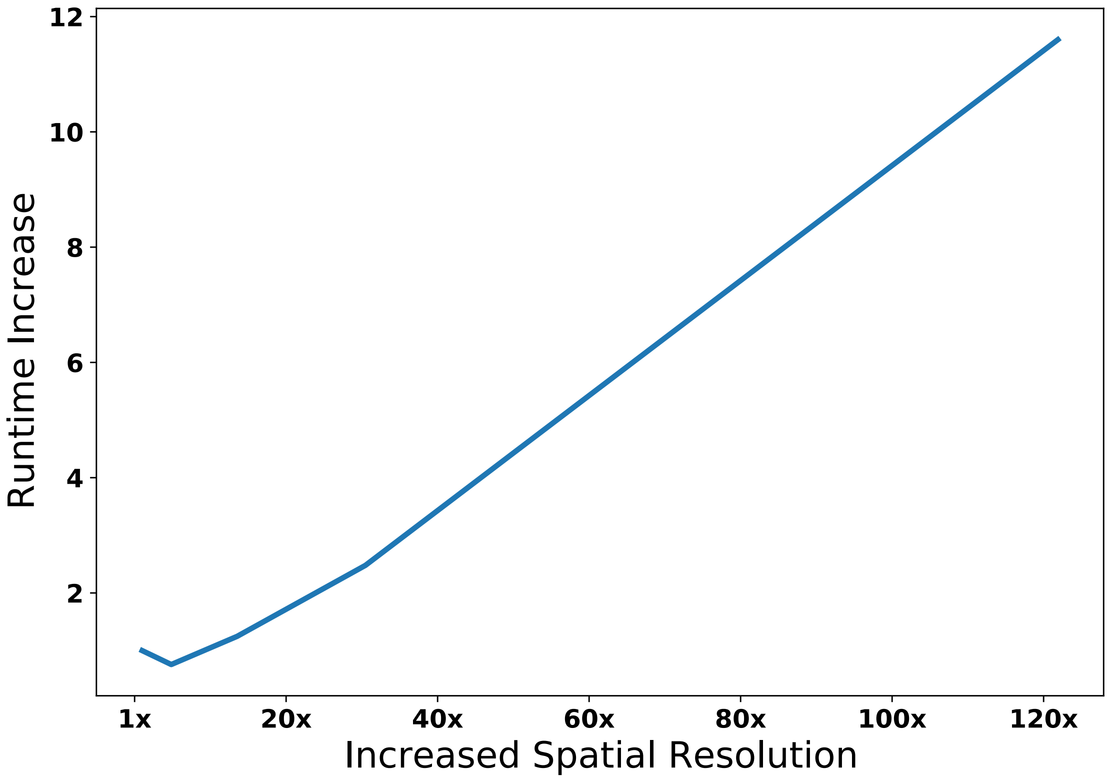


**Figure 9:** BayesTME runtime scalessub-linearly with the number of spots.

is the discrete spot deconvolution which scales quadratically with the number of possible cells in a given spot. However, as the resolution increases, this number actually decreases. The result is that the quadratic increase in spots is offset by the decrease in deconvolution burden. Figure 9 shows the relative runtime of BayesTME as we increase the spatial resolution from the current resolution (1x) with a few thousand nodes to high-resolution (120x) with hundreds of thousands of nodes. Despite increasing by two orders of magnitude, BayesTME only requires a 10x increase in computation time, which is tractable for modern compute clusters.

### 8 Posterior Sampling

**Sampling $R_{igk}$** Since $R_{igk} \sim Pois(\beta_k d_{ik} \phi_{kg})$,

$$R_{ig} = \sum_k R_{igk} \sim Pois(\sum \beta_k d_{ik} \phi_{kg}) \tag{1}$$

by the relationship between Poisson and Multinomial distribution, letting $\xi_{ik} = \frac{\beta_k d_{ik} \phi_{kg}}{\sum_k \beta_k d_{ik} \phi_{kg}}$, we can sample $R_{ig1}, \ldots, R_{igK}$ adjointly by

$$R_{ig1}, \ldots, R_{igK} | - \sim Mult(R_{ig}; \xi_{i1}, \ldots, \xi_{iK}) \tag{2}$$

**Sampling $\boldsymbol{\phi}_k$** Since $\boldsymbol{\phi}_k \sim Dir(\alpha_k)$, $\sum_g \phi_{kg} = 1$. One can show that

$$\begin{aligned} R_{ik1}, \ldots, R_{ikG} &\sim Mult(\sum_g R_{ikg}; \frac{\lambda_{ik}\phi_{k1}}{\sum_g \lambda_{ik}\phi_{kg}}, \ldots, \frac{\lambda_{ik}\phi_{kG}}{\sum_g \lambda_{ik}\phi_{kg}}) \\ &= Mult(\sum_g R_{ikg}; \phi_{k1}, \ldots, \phi_{kG}) \end{aligned} \tag{3}$$

Thus,

$$\boldsymbol{\phi}_k | - \sim Dir(\alpha_k + \sum_i R_{ik1}, \ldots, \alpha_k + \sum_i R_{ikG}) \tag{4}$$

**Sampling $\beta_k$** The likelihood of reads $R_{ik\cdot} = (R_{ik1}, \ldots, R_{ikG})$ is

$$\begin{aligned} P(R_{ik\cdot}|\beta_k, d_{ik}, \phi_{kg}) &= \prod_g P(R_{igk}|\beta_k, d_{ik}, \phi_{kg}) \\ &= \prod_g \frac{e^{-\beta_k d_{id} \phi_{kg}} (\beta_k d_{ik} \phi_{kg})^{R_{igk}}}{R_{igk}!} \\ &= \frac{e^{-\sum_g \beta_k d_{ik} \phi_{kg}} \prod_g (\beta_k d_{ik} \phi_{kg})^{R_{ikg}}}{\prod_g R_{ikg}!} \\ &= \frac{e^{-\beta_k d_{ik} \sum_g \phi_{kg}} (\beta_k d_{ik})^{\sum_g R_{ikg}} \prod_g (\phi_{kg}^{R_{ikg}})}{\prod_g R_{ikg}!} \\ &= \frac{e^{-\beta_k d_{ik}} (\beta_k d_{ik})^{R_{ik}} \prod_g (\phi_{kg}^{R_{ikg}})}{\prod_g R_{ikg}!} \\ &= \frac{\prod_g \phi_{kg}^{R_{ikg}}}{\prod_g R_{ikg}!} e^{-\beta_k d_{ik}} (\beta_k d_{ik})^{R_{ik}} \end{aligned} \tag{5}$$

and the posterior can be written as

$$\begin{aligned} P(\beta_k|R_{ikg}, d_{ik}, \phi_{kg}) &= \prod_i \prod_g P(R_{igk}|\beta_k, d_{ik}, \phi_{kg}) P(\beta_k) \\ &\propto \prod_i P(R_{ik\cdot}|\beta_k, d_{ik}, \phi_{kg}) P(\beta_k) \\ &= \prod_i \left( \frac{\prod_g \phi_{kg}^{R_{ikg}}}{\prod_g R_{ikg}!} e^{-\beta_k d_{ik}} (\beta_k d_{ik})^{R_{ik}} \right) P(\beta_k) \\ &= \left( \prod_i \frac{\prod_g \phi_{kg}^{R_{ikg}}}{\prod_g R_{ikg}!} \right) e^{-\beta_k \sum_i d_{ik}} \prod_i (\beta_k d_{ik})^{R_{ik}} P(\beta_k) \\ &= \left( \prod_i \frac{\prod_g \phi_{kg}^{R_{ikg}}}{\prod_g R_{ikg}!} \right) e^{-\beta_k \sum_i d_{ik}} \beta_k^{\sum_i R_{ik}} \prod_i d_{ik}^{R_{ik}} P(\beta_k) \\ &= \left( \prod_i \frac{d_{ik}^{R_{ik}} \prod_g \phi_{kg}^{R_{ikg}}}{\prod_g R_{ikg}!} \right) e^{-\beta_k \sum_i d_{ik}} \beta_k^{\sum_i R_{ik}} \frac{b^a}{\Gamma(a)} \beta_k^{a-1} e^{-b\beta_k} \\ &\propto e^{-\beta_k \sum_i d_{ki}} \beta_k^{\sum_i R_{ik}} \beta_k^{a-1} e^{-b\beta_k} \\ &= \beta_k^{\sum_i R_{ik} + a - 1} e^{-\beta_k (\sum_i d_{ki} + b)} \end{aligned} \tag{6}$$

Thus,

$$\beta_k|- \sim Gamma(a + \sum_i R_{ik}, b + \sum_i d_{ki}) \tag{7}$$

**Sampling $D_i$ and $\boldsymbol{d}_i$** The posterior distribution of the cell number of each type can be formulated as

$$P(d_{ik}|\boldsymbol{R}_i, \boldsymbol{\phi}_k, \beta_k, \boldsymbol{\psi}_i) = P(\boldsymbol{R}_{i,k:K}|d_{ik}, \boldsymbol{\theta}_i, \boldsymbol{r})P(d_{ik}, R_{i,1:k-1}) \tag{8}$$

We can be model the cell number distribution by a HMM type model, where we have $l_k = (d_k, n_k)$ as the latent stats, and $\boldsymbol{R}_{ik}$ as the observation. With a forward-filtering backward-sampling style algorithm, we can

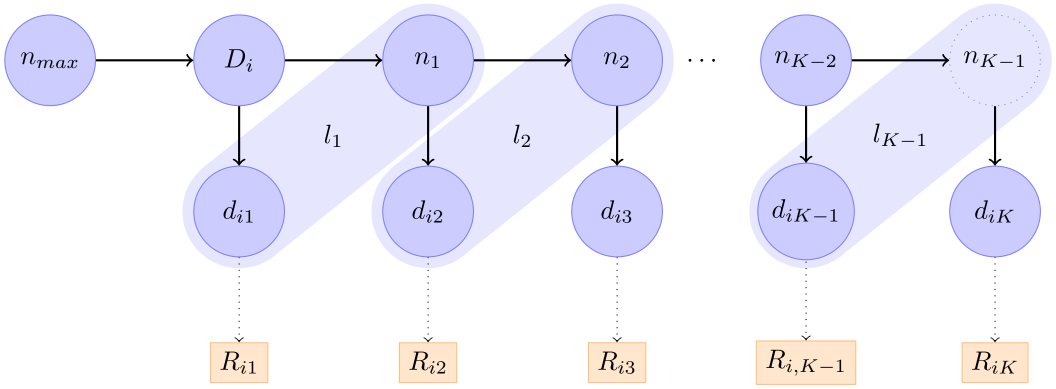


**Figure 10:** HMM modeling of cell numbers

calculate

$$\begin{aligned}
\alpha(l_k) &= P(l_k, \boldsymbol{R}_{1:k}) \\
&= P(R_k|l_k) \sum_{l_{k-1}} P(l_k|l_{k-1})\alpha(l_{k-1}) \\
&= P(R_k|d_k, n_k) \sum_{n_{k-1}} \sum_{d_{k-1}} P(l_k|d_{k-1}, n_{k-1})\alpha(d_{k-1}, n_{k-1}) \\
&= P(R_k|d_k) \sum_{n_{k-1}} \sum_{d_{k-1}} P(l_k|n_{k-1})\alpha(d_{k-1}, n_{k-1}) \quad * \\
&= P(R_k|d_k) \sum_{n_{k-1}} \left( P(l_k|n_{k-1}) \sum_{d_{k-1}} \alpha(d_{k-1}, n_{k-1}) \right) \\
&= P(R_k|d_k)P(l_k|n_{k-1}) \sum_{d_{k-1}} \alpha(d_{k-1}, n_{k-1}) \quad \star
\end{aligned} \tag{9}$$

and

$$\begin{aligned}
\alpha(l_1) &= P(R_1|l_1) \sum_D P(l_1|D)P(D|n_{max}) \quad \star\star \\
&= P(R_1|d_1)P(l_1|D)P(D|n_{max})
\end{aligned} \tag{10}$$

where step $*$ follows $R_k$ is independent from $n_k$ and $l_k$ is independent from $d_{k-1}$, and step $\star$ follows that a specific latent variable pair $l_k = (d_k, n_k)$ can only be sampled from corresponding $n_{k-1} = d_k + n_k$, and similar reasons holds for $\star\star$.

Thus, we can sample from the cell number from the posterior

$$
\begin{aligned}
P(d_{ik}|l_{k+1}, \boldsymbol{R}_{1:T}) &= P(d_{ik}|n_k, l_{k+1}, \boldsymbol{R}_{1:K}) \\
&= \frac{P(d_{ik}, n_k|l_{k+1}, \boldsymbol{R}_{1:K})}{P(n_k|l_{k+1}, \boldsymbol{R}_{1:K})} \\
&\propto P(l_k|l_{k+1}, \boldsymbol{R}_{1:T}) \quad \dagger \\
&\propto P(l_k, l_{k+1}|\boldsymbol{R}_{1:T}) \\
&\propto \alpha(l_k)P(l_{k+1}|l_k) \quad \ddagger \\
&= \alpha(l_k)P(l_{k+1}|n_k)
\end{aligned}
\tag{11}
$$

and

$$
\begin{aligned}
P(d_{iK}|\boldsymbol{R}_{1:K}) &= P(n_{K-1}|\boldsymbol{R}_{1:K}) \\
&\propto P(n_{K-1}, \boldsymbol{R}_{1:K}) \\
&= P(\boldsymbol{R}_K|n_{K-1}, R_{1:K-1})P(n_{K-1}, R_{1:K-1}) \\
&= P(\boldsymbol{R}_K|n_{K-1}) \sum_{d_{iK-1}} \alpha(l_{K-1}) \\
P(d_{iK-1}|d_{iK}, \boldsymbol{R}_{1:K}) &= P(d_{iK-1}|n_{K-1}, \boldsymbol{R}_{1:K}) \\
&\propto P(d_{iK-1}, n_{K-1}, \boldsymbol{R}_{1:K}) \\
&= \alpha(l_{K-1})
\end{aligned}
\tag{12}
$$

where † follow $n_k$ is deterministic given $l_{k+1} = (d_{k+1}, l_{k+1})$ and details for ‡ is shown in Equation (13)

$$
\begin{aligned}
P(l_k, l_{k+1}|R_{1:K}) &= P(R_{1:k}, R_{k+1}, R_{k+2:K}, l_{k+1}, l_k) \\
&= P(R_{k+2:K}|R_{1:k}, R_{k+1}, l_{k+1}, l_k)P(R_{1:k}, R_{k+1}, l_{k+1}, l_k) \\
&= P(R_{k+2:K}|l_{k+1})P(R_{k+1}|R_{k+1}, l_{k+1}, l_k)P(R_{1:k}, l_{k+1}, l_k) \\
&= P(R_{k+2:K}|l_{k+1})P(R_{k+1}|l_{k+1})P(l_{k+1}|R_{1:k}, l_k)P(R_{1:k}, l_k) \\
&= P(R_{k+2:K}|l_{k+1})P(R_{k+1}|l_{k+1})P(l_{k+1}|l_k)P(R_{1:k}, l_k) \\
&= \beta(l_{k+1})P(R_{k+1}|l_{k+1})P(l_{k+1}|l_k)\alpha(l_k) \\
&\propto \alpha(l_k)P(l_{k+1}|l_k)
\end{aligned}
\tag{13}
$$

**Spatial Smoothing via GFBT.** We proposed a graph fused binomial tree (GFBT) model to perform the spatial smoothing on cell assignment probabilities. Based on the result from HMM, we can construct a binomail tree model as in Figure 11. Let $p_{ki}$ denote the probability that a cell from cell type $k$ is observed in patch $i$. In the tree model, we break such cell assignment probability into a series of binomial decisions. The black nodes indicate the cell types and red nodes are the binomial decision nodes with $\varphi_k$ probability going to the left and $1 - \varphi_k$ probability going to the right, and $r_k = \sum_{j=k}^{K} d_{ji}$.

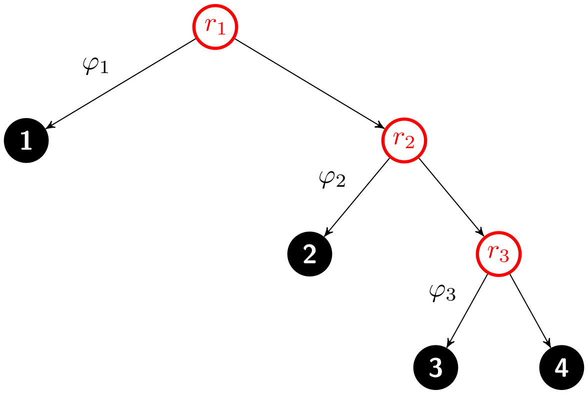


**Figure 11:** Example of the binomial tree with 4 type of cells

Let $\varphi_k = \sigma(\theta_{ki})$, we have $d_{ki} \sim Bin(r_k, \sigma(\theta_{ki}))$. Thus,

$$\begin{aligned}
d_{ki} &\sim Bin\left(r_k, \frac{e^{\theta_{ki}}}{1+e^{\theta_{ki}}}\right) \\
p(d_{ki}|\theta_{ki}) &= \binom{r_k}{d_{ki}} \left(\frac{e^{p_{ki}}}{1+e^{p_{ki}}}\right)^{d_{ki}} \left(\frac{1}{1+e^{p_{ki}}}\right)^{r_k - d_{ki}} \\
&\propto \frac{(e^{p_{ki}})^{d_{ki}}}{(1+e^{p_{ki}})^{r_k}} \\
&= 2^{-r_k} e^{\kappa_{ki}\theta_{ki}} \int_0^\infty e^{\omega_{ki}\theta_{ki}^2/2} p(\omega) d\omega \\
&\propto e^{\kappa_{ki}\theta_{ki}} \mathbb{E}_\omega \left[e^{\omega p_{ki}^2/2}\right] \\
p(\boldsymbol{\theta}|\cdot) &\propto p(\boldsymbol{\theta}) p(\mathbf{d}|\boldsymbol{\theta}, \cdot) \\
&= p(\boldsymbol{\theta}) \prod_{k,i} \exp\left[\kappa_{ki}\theta_{ki} - \frac{\omega_{ki}}{2}\theta_{ki}^2\right] \\
&= p(\boldsymbol{\theta}) \exp\left[-\frac{1}{2}(\mathrm{vec}(\boldsymbol{\theta}) - \mathrm{vec}(\mathbf{z}))^T \Omega (\mathrm{vec}(\boldsymbol{\theta}) - \mathrm{vec}(\mathbf{z}))\right]
\end{aligned} \tag{14}$$

where

$$\begin{aligned}
\Omega &= \mathrm{diag}(\omega_{11}, \ldots, \omega_{KN}) \\
\omega_{ki} &\sim PG(r_k, \theta_{ki}) \\
z_{ki} &= \frac{\kappa_{ki}}{\omega_{ki}} \\
\kappa_{ki} &= d_{ki} - \frac{r_k}{2}
\end{aligned} \tag{15}$$

The prior on $\Delta\theta_{ki}$ is $\mathcal{N}(0, \lambda_{ki}^2\tau_{ki}^2)$, then with $\Psi = \Delta^T \mathcal{T} \Delta \otimes I_K$, the exponent term in the pdf function is

$$\begin{aligned}
&(\boldsymbol{\theta} - \mathbf{z})^T \Omega (\boldsymbol{\theta} - \mathbf{z}) + (\boldsymbol{\theta} - \mathbf{z})^T \Psi (\boldsymbol{\theta} - \mathbf{z}) \\
=&\boldsymbol{\theta}^T (\Omega + \Psi) \boldsymbol{\theta} - 2(\mathbf{z}^T \Omega) \boldsymbol{\theta} + \boldsymbol{\theta}^T \Omega \boldsymbol{\theta} \\
\propto&\boldsymbol{\theta}^T \Sigma \mathbf{p} - 2\boldsymbol{\mu}^T \boldsymbol{\theta} \\
\propto&(\boldsymbol{\theta} - \Sigma^{-1} m) \Sigma (\boldsymbol{\theta} - \Sigma^{-1} m)
\end{aligned} \tag{16}$$

where $\mathbf{p}$ and $\mathbf{z}$ refer to the vectorized $\mathbf{p}$ and $\mathbf{z}$

$$\begin{aligned} \Sigma &= \Omega + \Psi \\ m &= \Omega \mathbf{z} \\ &= \Omega \Omega^{-1} \kappa \\ &= \kappa \end{aligned} \tag{17}$$

Thus, we have

$$\begin{aligned} \boldsymbol{\theta} | \cdot &\sim \mathcal{N}(\mu, \Sigma^{-1}) \\ \Sigma &= \Omega + \Delta^T \mathcal{T} \Delta \otimes I_K \\ \mu &= \Sigma^{-1} \kappa \end{aligned} \tag{18}$$

we have the resampling scheme

$$\begin{aligned} \mathcal{T} &= \text{diag}(\frac{1}{\lambda^2 \tau^2}) \\ \omega_{ki} &\sim PG(r_k, \theta_{ki}) \\ \Omega &= \text{diag}(\omega_{11}, \ldots, \omega_{KN}) \\ \Sigma^{-1} &= \Delta^T \mathcal{T} \Delta \otimes I_K + \Omega \\ \text{vec}(\boldsymbol{\theta}) &\sim \mathcal{N}(\Sigma^{-1} \text{vec}(\kappa), \Sigma^{-1}) \end{aligned} \tag{19}$$

With $\theta$ after spatial smoothing, we can recover the cell assignment probability

$$p_k = \begin{cases} \sigma(\theta_{ki}) \prod_{j=1}^{k-1} (1 - \sigma(\theta_{ji})) & k < K \\ \prod_{j=1}^{K} (1 - \sigma(\theta_{ji})) & k = K \end{cases} \tag{20}$$

where $K$ is the total number of cell types.
